# Endosomal pH Triggers Amyloid β Oligomerization and Maladaptive Phenotypic Plasticity in Alzheimer’s Disease

**DOI:** 10.64898/2026.09.23.752284

**Authors:** Ritesh Kumar Meena, Atchuta Srinivas Duddu, Mohit Kumar Jolly, Hari Prasad

**Affiliations:** Centre for Brain Research, Indian Institute of Science Campus, Bengaluru, 560012, India; Department of Bioengineering, Indian Institute of Science, Bengaluru, 560012, India

**Author notes:** Address for correspondence: Dr. Hari Prasad, MBBS, MMST, PhD, Centre for Brain Research, Indian Institute of Science Campus, Bengaluru, 560012, India,. Equal contribution (Ritesh Kumar Meena, Atchuta Srinivas Duddu).

**Keywords:** Endosomal pH, NHE6, APOE4, Amyloid β, Phenotypic plasticity, Alzheimer’s disease. Running title: Endosomal pH Promotes Aβ Oligomers and Dysregulated Plasticity

## Abstract

Endosomal dysfunction is a presymptomatic hallmark of neurodegeneration. Recent evidence highlights dysregulation of endosomal pH as a central pathogenic hub in Alzheimer’s disease (AD); however, the mechanisms linking pH shifts to neurodegeneration remain incompletely defined. Here, we use a quantitative model of endosomal acidification driven by proton pumping via the vacuolar ATPase, proton leak via the endosomal Na /H exchanger NHE6, and other ion-regulating elements. The model recapitulates how downregulation of NHE6 in AD promotes endosomal hyperacidification, potentially triggering maladaptive phenotypic plasticity—an initially adaptive response that becomes pathological. Analysis of human brain datasets reveals reciprocal enrichment of NHE6 in neurons and the related NHE9 in glia, with NHE6 co-expression networks enriched for synaptic signalling. Systematic curation of NHE6 patient variants indicates that loss-of-function is associated with late regression, consistent with progressive endosomal hyperacidification, supporting a conceptual framework where early compensation transitions to neurodegeneration. Mathematical analyses calibrated for neuronal endosomes reveal a saturable relationship between luminal pH and NHE6 dosage, with threshold-like behaviour below ∼50% expression that hyperacidifies endosomes, correlating with AD severity. Our model suggests this pH shift may exponentially accelerate Aβ oligomerization and enhance β-secretase activity. Furthermore, Aβ oligomerization estimates correlate with dysregulation of calcium signalling and synaptic dysfunction. Model findings are compared with experimental results from NHE6-null mice and a cell culture model of AD. Drawing parallels to cancer, we propose that endosomal pH serves as a conserved regulator of adaptive-to-maladaptive transitions. Restoring physiological endosomal pH may offer a therapeutic window to prevent irreversible neurodegeneration in AD.

## Introduction

Alzheimer’s disease (AD) is a progressive neurodegenerative disorder characterized by synaptic failure, cognitive decline, and the accumulation of amyloid-β (Aβ) plaques and neurofibrillary tangles^1^. At the clinicopathological level, increasing evidence suggests that dysfunction of early endosomes precedes overt pathology by decades^2,3^. An emerging “hub-and-spoke” model positions endosomal function as a central pathogenic hub, from which Aβ, tau, synaptic, and microglial pathologies radiate, highlighting endosomal defects as a potential initiating event in AD pathogenesis^4^. Despite nearly three decades of research, the underlying molecular mechanisms and cellular consequences of endosomal dysfunction remain incompletely understood, and biomarkers or disease-modifying therapies targeting endosomes are still lacking.

A central feature of endosomal biology is luminal pH regulation, which governs vesicle trafficking. Endosomal acidification is tightly regulated by proton pumping via vacuolar ATPase (V-ATPase), counterbalanced by proton leak pathways mediated by endosomal Na /H exchangers (eNHEs), namely NHE6 (*SLC9A6*) and NHE9 (*SLC9A9*)^5–8^. These exchangers establish a dynamic pH setpoint critical for proper endosomal function. This proton pump–leak system is evolutionarily conserved from yeast to plants and mammals, underscoring the essential role of luminal pH in vesicular trafficking^9^. Though these exchangers are ubiquitously expressed, they are particularly enriched in the brain^10^. Disruption of eNHE function is increasingly recognized across neurodevelopmental, psychiatric, and neurodegenerative disorders, suggesting a shared pathogenic mechanism^11^. Loss-of-function mutations in NHE6 cause Christianson syndrome (CS), a severe X-linked disorder characterized by developmental delay, intellectual disability, absent speech, seizures, and ataxia^12^.

Maintaining ion and pH homeostasis—especially within the endosomal system—is not just a cellular housekeeping task but a fundamental determinant of long-term brain health^9^. These pH and ion dynamics operate on fast time scales (milliseconds) in comparison to other biological processes including metabolic processes (hours), immune shifts (days), and the development of plaques and atrophy (years)^13^. In AD, pathology does not arise from a single failure but rather from mismatches across these rhythms, where short-term disruptions accumulate and lead to long-term neurodegeneration^13^. Accordingly, even subtle, chronic dysregulation of endosomal pH could initiate a cascade of mismatched events, ultimately driving the protracted neurodegenerative course of AD. Among regulators of endosomal pH, NHE6 has emerged as a critical molecular node linked to AD pathogenesis. Previous work demonstrated that NHE6 transcript and protein levels are reduced in AD and inversely correlate with disease severity^14^. Additional studies have identified NHE6 as a key player in early-stage AD^15^, a central hub with 202 network connections in AD^16^, and a major regulator of trafficking and proteostatic clearance pathways implicated in AD^17^. Deep-learning approaches have also highlighted NHE6 as a predictive feature for AD^18^. Moreover, transcriptomic analysis shows that NHE6 is downregulated (up to sixfold) in older (70 years) compared to younger (40 years) brain^19^. Importantly, the ε4 allele of apolipoprotein E (APOE4), the strongest genetic risk factor for sporadic AD, is associated with reduced NHE6 expression in both post-mortem brain tissue and cellular models^20,21^.

While endosomal hyperacidification is still emerging as a potential mechanism in AD, lysosomal hypoacidification is well documented and is associated with reduced activation of pH-sensitive cathepsins and impaired autophagy^22^. However, some evidence also points to lysosomal hyperacidification, accompanied by increased cathepsin activation in the context of AD, which may contribute to autophagic dysfunction, albeit through a distinct mechanism^23^. Of note, observations in APOE4 astrocytes reveal endosomal hyperacidification linked to NHE6 downregulation, along with lysosomal hypoacidification associated with V-ATPase downregulation, indicating that these pH abnormalities can coexist^20^. The complexity of pH dysregulation is also evident in Parkinson’s disease (PD), where both lysosomal hypoacidification and hyperacidification have been linked to pathology, underscoring that lysosomal function requires an optimal “Goldilocks” pH range^24^. Together, these observations support a model in which pH extremes in either direction disrupt endolysosomal homeostasis, with loss of NHE6 resulting in hyperacidification of early endosomes that may contribute to secondary lysosomal dysfunction^25^.

Amyloidogenic processing of amyloid precursor protein (APP) is sensitive to endosomal pH, as β-secretase BACE (β-site amyloid precursor protein cleaving enzyme) activity has an acidic pH optimum^26^. Emerging evidence also indicates that acidic pH regulates Aβ aggregation kinetics and oligomer formation^27^. Despite this, experimental studies have shown that the role of NHE6 function and endosomal acidification in AD may not be straightforward. In some contexts, NHE6 deficiency corrects APOE4-associated trafficking defects and reduces plaque burden^28,29^, whereas in others it increases Aβ production and contributes to APOE4-associated reduced Aβ clearance in astrocytes^14,20,30^. Furthermore, ketamine, an anaesthetic and addictive drug, has been shown to promote the amyloidogenic pathway by downregulating NHE6 in mouse models^31^. Conversely, more recent studies activating the NHE6-endosomal pH axis have restored Aβ clearance and cognitive function in AD mouse models^32^. This raises a critical question: How can NHE6 downregulation be both protective and pathogenic?

These seemingly divergent results can be unified by drawing insights from cancer research, framing NHE6 downregulation as a promoter of maladaptive phenotypic plasticity—an initially adaptive cellular response to specific insults that, when sustained, becomes the engine of pathology^33–36^. For example, in colorectal cancer, endosomal pH alkalinization via upregulation of related NHE9 triggers a starvation response that promotes cell survival but ultimately leads to epithelial–mesenchymal transition and metastasis^37,38^. Similarly, NHE6 loss may initially mitigate some AD-related defects, but chronic downregulation of NHE6 may push endosomal pH past a critical threshold, turning this compensatory response into a maladaptive cascade. In this study, we integrate single-cell transcriptomics, human genetic variant analysis, pathway enrichment, and quantitative mathematical modeling of endosomal pH dynamics in the context of AD. By linking NHE6 dosage to endosomal acidification, Aβ production and oligomerization kinetics, we define a mechanistic continuum that connects luminal pH dysregulation to disease-relevant phenotypes, including Aβ oligomerization, and neurodegeneration. More broadly, this work supports a model in which dysregulation of endosomal Na /H exchangers promote maladaptive phenotypic plasticity, providing a unified framework linking dysfunction of the early endosomal compartment to both neurodegeneration and cancer.

## Results

### Reciprocal Enrichment of Endosomal Na /H Exchangers in Neurons and Glia Across Human Brains

To establish the cellular landscape of eNHEs in the human brain, we first examined single-cell transcriptomic data from two independent normal human brain datasets: LIBD and IsoHuB. Across both datasets, we observed distinct expression patterns: NHE6 was highly expressed in neurons, including both excitatory and inhibitory subtypes, whereas NHE9 was predominantly expressed in glial cells, such as microglia and oligodendrocyte precursor cells (*Fig. 1A-D and Supplementary Fig. 1A-B*). This reciprocal enrichment suggests functional specialization of these isoforms, offering a potential explanation for their nonredundant roles and the distinct clinical phenotypes associated with their genetic aberrations, despite their overlapping subcellular localization in endosomes^10^. A similar expression pattern was observed across six additional single-cell datasets from various brain pathologies curated by the PsychENCODE Consortium (*Supplementary Fig. 1A*). Thus, in both normal and diseased human brains, eNHE expression exhibits a reciprocal neuronal-glial distribution.

**Figure 1:**
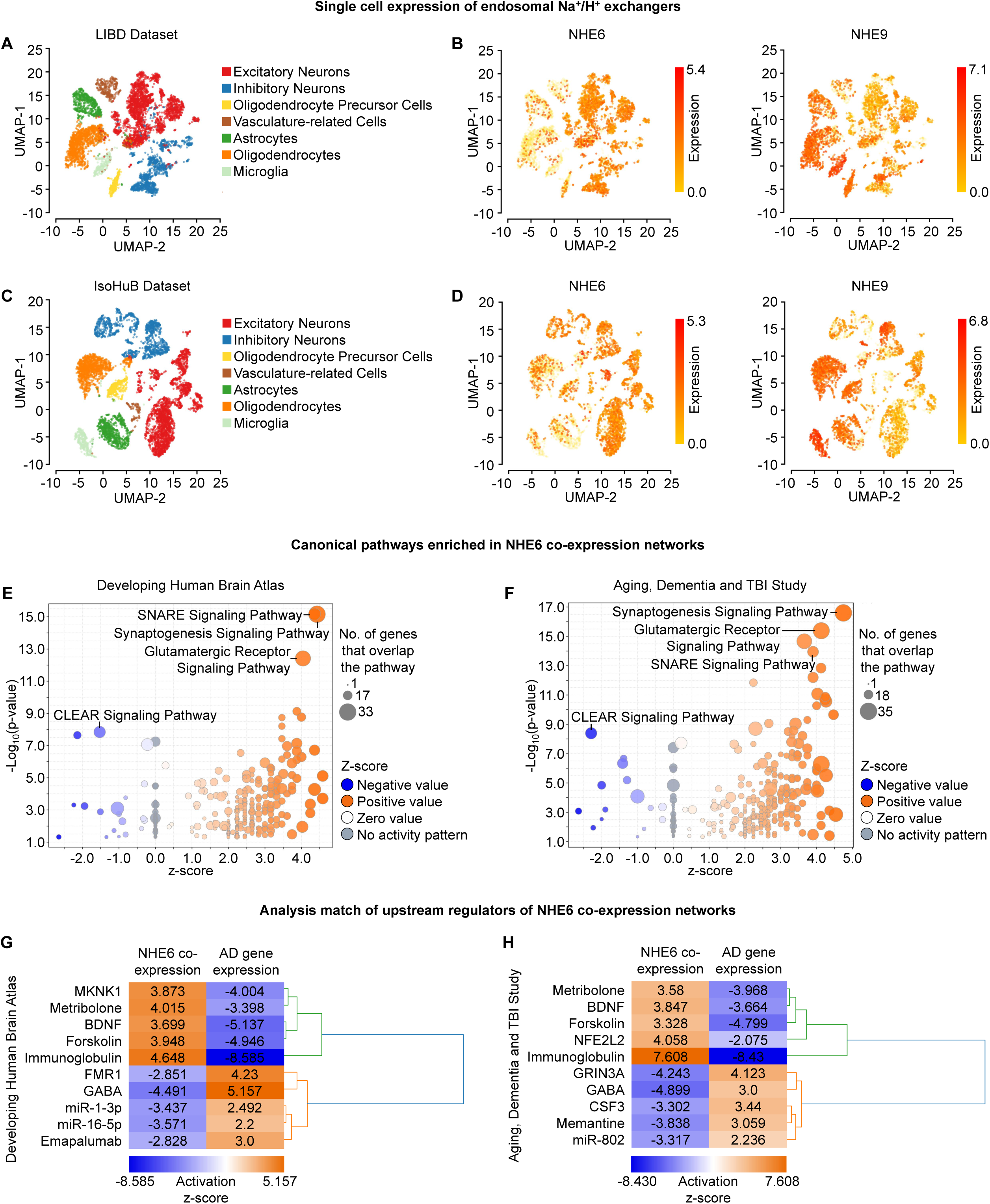
Single-cell analysis and functional network of NHE6 expression in the human brain. (A-D) Single-cell transcriptomic analysis of NHE6 expression compared with NHE9 in two independent normal human brain datasets: LIBD (n=4) (A, B) and IsoHuB (n=10) (C, D). UMAP plots show cell type clusters (A, C) and cell-type-specific expression patterns (B, D). Note that NHE6 is enriched in neuronal populations, including excitatory and inhibitory neurons, whereas NHE9 is predominantly expressed in glial populations, including microglia and oligodendrocyte precursor cells. Color intensity indicates normalized expression levels. (E, F) Volcano plots depicting canonical pathways enriched in NHE6 co-expression networks identified by Ingenuity Pathway Analysis (IPA) of the top 500 genes co-expressed with NHE6 in the Developing Human Brain Atlas (E) and the Aging, Dementia, and TBI Study (F) datasets. Circle size indicates the number of genes overlapping with each pathway, and color represents activation z-scores (red: activated, blue: inhibited). The top three significantly enriched canonical pathways include SNARE signaling, synaptogenesis signaling, and glutamatergic receptor signaling; the top inhibited canonical pathway is the CLEAR (Coordinated Lysosomal Expression and Regulation) pathway. (G, H) Heatmaps of the Analysis Match comparison of upstream regulator activation states between NHE6 co-expression signatures and the IPA knowledge base reveal opposing patterns with AD postmortem brain datasets (GSE129308, G; GSE36980, H) (red: activated, blue: inhibited). Note the reciprocal regulation of forskolin and BDNF—activated in NHE6 co-expression signatures but inhibited in AD—consistent with CREB-mediated regulation of NHE6 expression. See also Supplementary Figure 1.

To further explore the functional context of NHE6, we leveraged the principle that genes that express together often function together in shared biological pathways. Accordingly, we identified the top 500 genes co-expressed with NHE6 using gene–gene co-expression matrices derived from the Developing Human Brain Atlas and the Aging, Dementia, and TBI Study RNA-seq datasets, followed by Ingenuity Pathway Analysis (IPA) (*Supplementary Fig. 1C*). Canonical pathways enriched among NHE6-coexpressed genes included SNARE (soluble-ethylmaleimide-sensitive factor attachment protein receptor) signalling, synaptogenesis signalling, and glutamatergic receptor signalling, highlighting a strong association with synaptic function (*Fig. 1E-F*). The identification of SNARE signalling is especially noteworthy (*Supplementary Fig. 1D*), given previous work implicating NHE6 in this pathway in the context of PD^39^.

We next employed the Analysis Match function in IPA to compare upstream regulators of NHE6-coexpressed genes (*Supplementary Fig. 1E*) with disease-associated gene expression datasets. Notably, AD emerged as a condition showing a negative association with NHE6 co-expression signatures across datasets, suggesting that downregulation of NHE6 may contribute to AD pathophysiology, as we previously reported^14,20^ (*Fig. 1G-H*). Strikingly, several upstream regulators exhibited reciprocal regulation trends between NHE6-coexpressed genes and postmortem AD brains. In particular, forskolin and BDNF (brain-derived neurotrophic factor)—both associated with activation of cAMP signalling— were activated in NHE6 co-expression signatures, consistent with experimental evidence that NHE6 is a target of the transcription factor cAMP-response element-binding protein (CREB)^40^ (*Fig. 1G-H and Supplementary Fig. 1E*). Conversely, both upstream regulators were found to be inhibited in AD brains (*Fig. 1G-H*). Together, these findings highlight a neuron-enriched NHE6 regulatory network that is disrupted in AD, setting the stage for exploring how endosomal pH dysregulation may promote maladaptive phenotypic plasticity.

### Genetic Variants of NHE6 Reveal a Link Between Loss of Function and Maladaptive Plasticity

Mutations in NHE6 provide crucial insight into the dominant mechanisms driving neurological disorders linked to NHE6 dysfunction. Much of the existing literature is limited to single case reports, and a comprehensive overview has been lacking. To address this, we performed a systematic literature survey and compiled a curated list of 120 NHE6 patient variants associated with a range of phenotypes (*Supplementary Table 1*). These variants span a gamut of mutation types: small deletions and missense substitutions are the most common, followed by splicing variants, nonsense mutations, small insertions, and gross deletions. Regulatory, start-loss, and small indel variants are rare, with one, two, and three reported occurrences, respectively (*Fig. 2A and Supplementary Table 1*). Mapping missense and nonsense variants onto the NHE6 protein structure revealed that these lesions occur throughout the coding sequence, including the membrane-embedded ion transport domain and the C-terminal cytoplasmic tail (*Fig. 2B-C and Supplementary Fig. 2A*). Functional evaluation has been performed for only a subset of missense variants, leaving the majority uncharacterized^41^ (*Supplementary Fig. 2B and Supplementary Table 1*). Most of missense variants involve evolutionarily conserved residues and are likely pathogenic, whereas mutations at poorly conserved sites may represent benign polymorphisms or have subtle functional effects (*Supplementary Table 1*).

**Figure 2:**
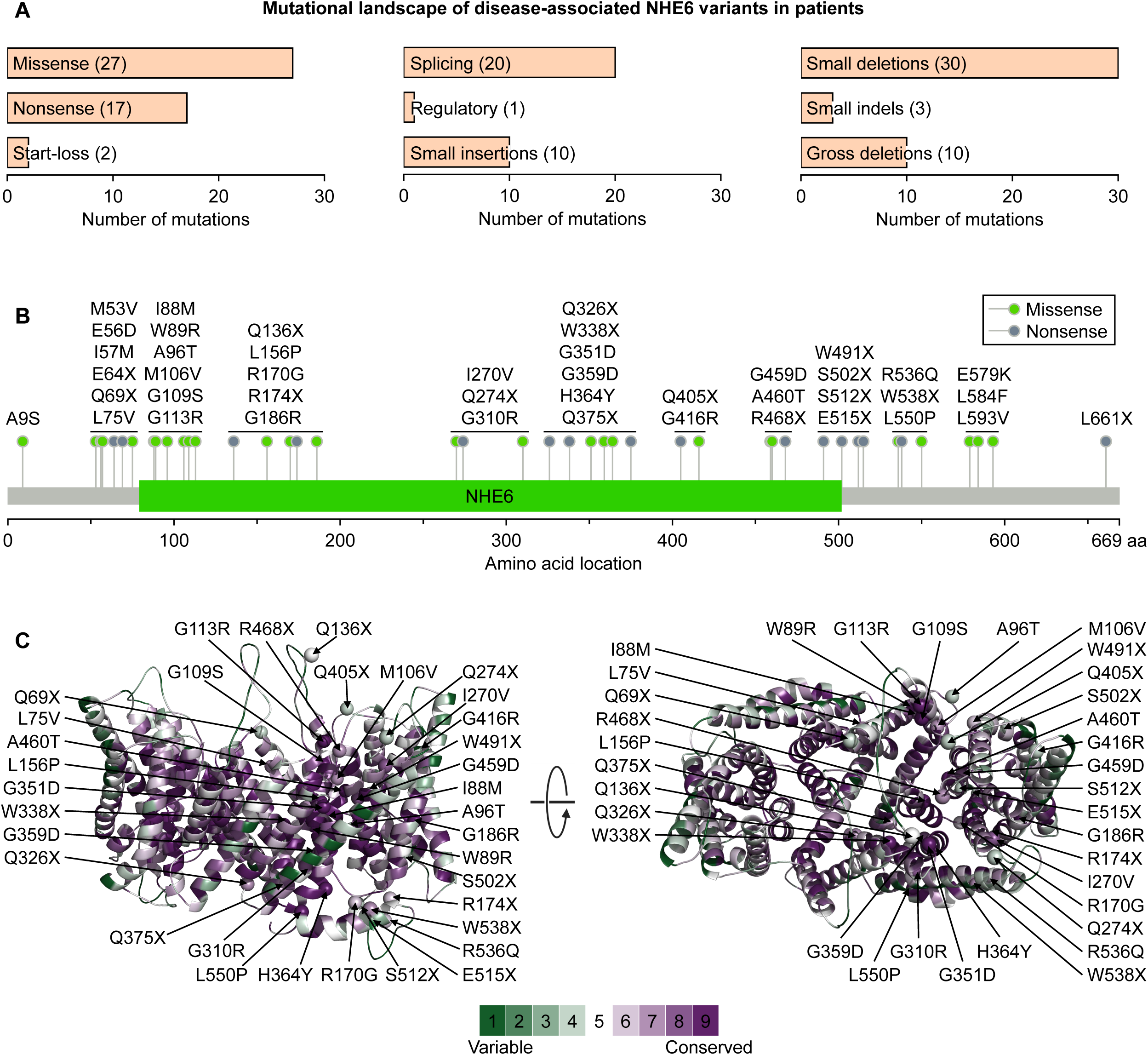
Systematic analysis of NHE6 patient variants associated with diverse disease phenotypes. (A) Bar graph depicting the distribution of 120 curated NHE6 patient variants by mutation type. Note that small deletions and missense variants represent the most frequent categories, followed by splicing variants, nonsense mutations, small insertions, and gross deletions. Rare categories include regulatory, start-loss, and small indel variants. The number in parentheses indicates the count of each mutation type. Predicted loss-of-function mutations constitute the predominant class. Disease phenotypes are provided in Supplementary Table 1. (B) Lollipop representation of NHE6 missense and nonsense variants mapped onto the coding sequence. The membrane-embedded ion transport domain is indicated. (C) Side (*left*) and top (*right*) views of the NHE6 dimer, including the transport domain and proximal C-terminal domain. The NHE6 model structure was generated based on the related NHE9 structure (PDB: 8PXB) and coloured according to evolutionary conservation scores, ranging from green (variable) to purple (highly conserved), using the SWISS-MODEL and ConSurf web servers. NHE6 missense and nonsense patient variants are depicted as α-carbon spheres on one monomer. The conservation colour bar is shown at the bottom. See also Supplementary Figure 2 and Supplementary Table 1.

Given that the majority of mutations are null variants—including splicing or truncating mutations that disrupt NHE6 function—loss of function appears to be a major pathogenic mechanism (*Supplementary Fig. 2B and Supplementary Table 1*). These null mutations are associated with severe phenotypes that cause CS, characterized by profound intellectual disability, cerebral atrophy, microcephaly, motor phenotypes, and a very early age of onset^12^. In contrast, missense mutations exhibit a broader phenotypic spectrum, ranging from mild forms, such as partial epilepsy without neurodevelopmental delay, to autism, and in some cases, severe phenotypes resembling those of null variants^12,42,43^. These observations suggest a dosage effect and provide a rationale for developing quantitative models to study how graded NHE6 levels or activity influence endosomal pH.

Another important consideration is the hallmark association of NHE6 loss of function with regression and progressive neurodegeneration^12^. Indeed, even in Dr. Christianson’s original description of an extended South African pedigree, early neurodevelopmental abnormalities were documented to be followed by a slow regression with age^44^. This supports the formulation that loss of NHE6 induces plasticity that can become maladaptive, leading to clinical regression and neurodegeneration. Notably, the only missense mutation associated with a gain of NHE6 function (p.G186R; NM_006359) is not associated with regression^45^ (*Supplementary Fig. 2B*). Together, these findings also point to a therapeutic window: early diagnosis and correction of endosomal pH defects resulting from loss of NHE6 function could prevent plasticity from becoming maladaptive before reaching a tipping point, thereby holding therapeutic potential. Of note, female carriers of NHE6 mutations exhibit a broad spectrum of outcomes, ranging from healthy to mild-to-moderate intellectual disability, psychiatric illness, autism, or PD-like features, and can also show regression^46,47^. This variability likely reflects X-chromosome mosaicism combined with dosage sensitivity, where the proportion of neurons expressing the mutant allele determines the severity and timing of phenotypic manifestations^46^. Thus, female carriers illustrate *in vivo* how graded NHE6 function can influence both early and progressive neurological outcomes.

These insights may also have implications for AD, where loss of NHE6 function has been reported to be associated with both beneficial and detrimental effects. At first glance, this duality appears inconsistent with patient mutations, in which loss of function is the predominant mechanism leading to severe phenotypes. However, this apparent contradiction can be reconciled by invoking a phenotypic plasticity framework, drawing insights from cancer biology, in which cellular responses to stress are initially adaptive but become maladaptive over time. Within this framework, downregulation of NHE6 and the resulting endosomal hyperacidification may trigger early cellular responses that are beneficial in specific contexts—for example, reactive microglia and astrocytes reducing amyloid plaque load or promoting endocytic recycling of certain receptors (e.g., the Reelin receptor ApoER2), as previously reported^28,29^— whereas prolonged hyperacidification-linked mechanisms might eventually cause maladaptation leading to neurodegeneration.

Support for this temporal model comes from post-mortem human CS brains carrying an in-frame deletion (p.W338_T340del; NM_006359), which show prominent glial activation^48^ (*Supplementary Fig. 2B*). Structure-function analyses of this mutation confirmed that it results in complete loss of NHE6 function^14,49^. This glial activation could be mediated through a primary cell-autonomous mechanism early in the disease course and may serve a compensatory role in limiting or delaying neurodegenerative pathology^50^. Clinically, patients harbouring this mutation also exhibit late-onset regression, which likely arises through secondary mechanisms induced by damaged neurons and tau deposition^48,51^. Neuroimaging of CS patients further supports this pathogenic model: neurodegenerative changes are less obvious early in life but become progressive in late childhood and adolescence, leading to severe cortical and cerebellar degeneration. Once maladaptive changes occur, they may accelerate neuronal loss and contribute to clinical regression^52^.

### Mathematical Model of NHE6 Downregulation and Endosomal pH in Alzheimer’s Disease

Having observed the neuronal enrichment of NHE6 and the dose-dependent phenotypic spectrum of its genetic variants, we next asked whether NHE6 downregulation in AD brains quantitatively predicts endosomal hyperacidification, and whether this hyperacidification exhibits a threshold behavior that could explain the transition from compensatory to maladaptive plasticity. To address this, we developed a coupled ODE-based mathematical model of endosomal pH regulation by adapting established frameworks for organellar acidification^53^ and incorporating NHE6-mediated proton extrusion. The model integrates the coordinated activity of V-ATPase (proton pumping), NHE6 (proton leak in exchange for Na ), Cl /H antiporter-mediated counterion flux (which dissipates the membrane potential generated by proton accumulation), and other relevant ion-regulating elements (*Fig. 3A*). We then parameterized the model for neuronal endosomes, using published experimental measurements and electrophysiological properties to calibrate transporter copy numbers, together with literature-derived values for endosomal physical parameters such as surface area, volume, and buffering capacity (*Fig. 3B–C and Supplementary Table 2*).

**Figure 3:**
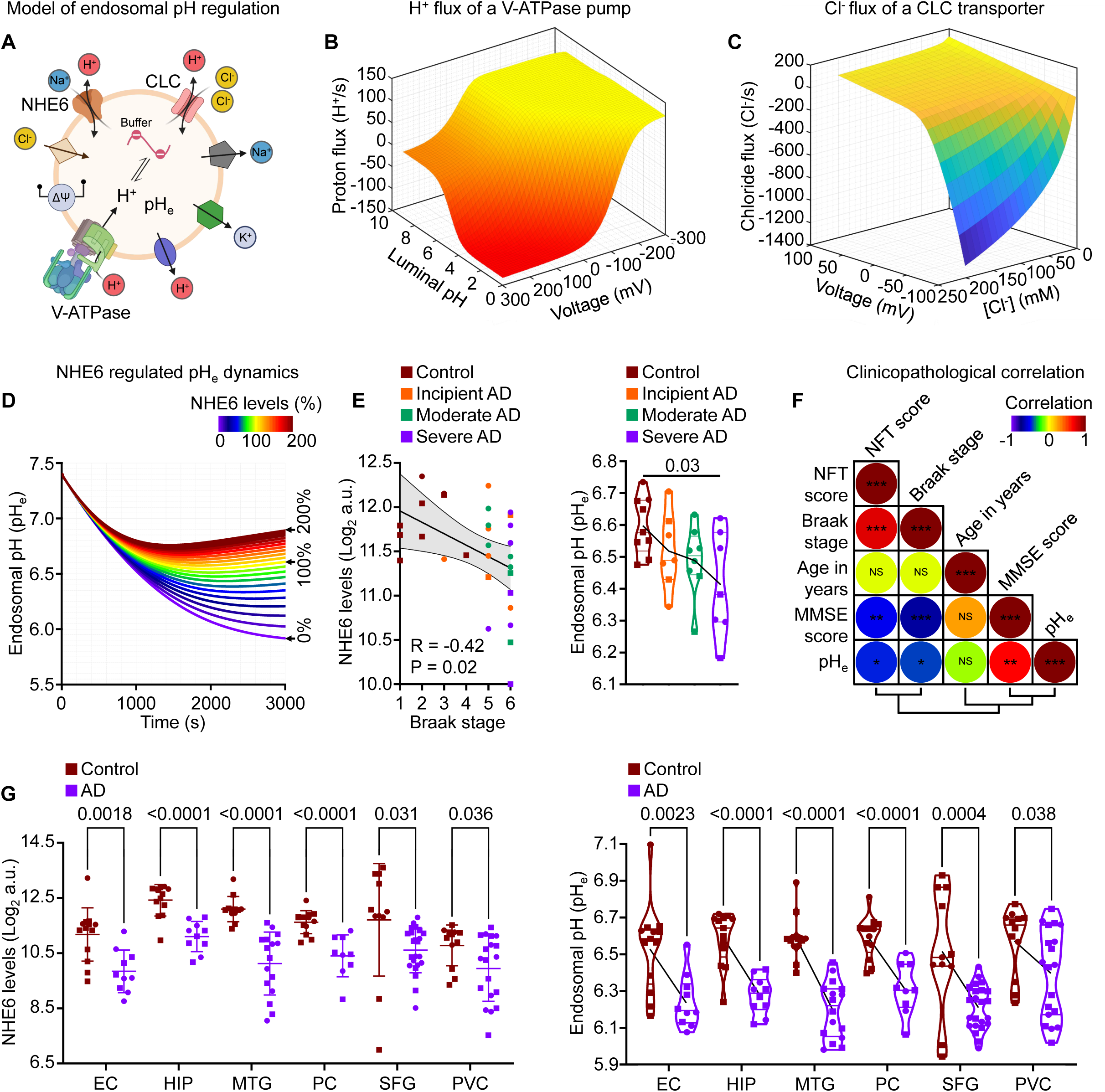
Quantitative modeling links NHE6 downregulation to endosomal hyperacidification in Alzheimer’s disease. (A) Schematic of the endosomal pH model integrating proton pumping via V-ATPase, proton leak mediated by NHE6, CLC Cl /H antiporter-mediated counterion flux, and additional ion-regulating mechanisms including passive leaks and buffering capacity controlling luminal acidification. The illustration was created with BioRender. (B, C) Proton pumping profile for V-ATPase (B) and chloride pumping profile for the CLC antiporter (C). The surface for flux is based on equations described in the Methods. (D) Dynamics of endosomal acidification as a function of NHE6 expression level (percentage of baseline, ranging from 0-200%). Note the saturable Michaelis-Menten-type relationship, showing threshold-like behaviour below ∼50% normal expression, where the curve transitions from steep pH dependence to a plateau. Loss of the proton leak pathway, as seen in CS-associated NHE6 mutations, causes endosomal hyperacidification. Conversely, increased proton leak, as seen in the gain-of-function mutant p.G186R, causes endosomal alkalinization. (E) Analysis of postmortem hippocampus (GSE1297) across disease stages (Control, Incipient AD, Moderate AD, Severe AD) showing (*left*) a scatter plot of NHE6 expression versus Braak stage with linear fit, Pearson correlation (R), and 95% confidence interval band, and (*right*) a violin plot with scatter of estimated endosomal pH derived from NHE6 expression using the mathematical model. A line connecting the average pH values is overlaid. P values were calculated by one-way ANOVA. Note significant endosomal hyperacidification in severe AD compared to control. (F) Heatmap of the correlation coefficient matrix depicting clinicopathological associations of estimated endosomal pH with age in years, antemortem cognitive function assessed by Mini-Mental State Examination (MMSE) score, and postmortem neuropathology assessed by neurofibrillary tangle (NFT) score and Braak stage. NS, not significant; *P < 0.05, **P < 0.01, ***P < 0.001 for each comparison. (G) Independent validation in GSE5281 dataset showing (*left*) scatter plots with mean and standard deviation for NHE6 expression in control and AD brains and (*right*) violin plots with scatter showing estimated endosomal pH across six brain regions: entorhinal cortex (EC), hippocampus (HIP), medial temporal gyrus (MTG), posterior cingulate (PC), superior frontal gyrus (SFG), and primary visual cortex (PVC). A line connecting the average pH values is overlaid. P values were calculated by unpaired, two tailed t test. Each scatter point in (E) and (G) represents an individual postmortem brain; squares represent males and circles represent females. See also Supplementary Figure 3 and Supplementary Table 2.

Sensitivity analysis was performed on buffering capacity (20–60 mM/pH unit; baseline: 40 mM/pH unit) and NHE6 turnover rate (1100–1900 ions/s; baseline: 1500 ions/s). While buffering capacity influenced the initial acidification rate, turnover rate primarily affected steady-state pH. Deviations from baseline were minimal-to-modest (<0.5% and <2%, respectively), supporting model robustness (*Supplementary Fig. 3A-B*). Importantly, we observed the relationship between NHE6 levels and endosomal pH is not linear; instead, it follows a saturable Michaelis–Menten–type relationship, exhibiting threshold-like behaviour below ∼50% of normal NHE6 expression (*Fig. 3D and Supplementary Fig. 3C*). The marginal gain in pH for every 50% increase in NHE6 levels (ΔpH/ΔNHE6) declines progressively (0.87 → 0.51 → 0.33 → 0.24) (*Supplementary Fig. 3D*). Although the inflection point is not sharp, this relationship recapitulates the graded severity of patient mutations and may define a potential therapeutic window.

Incorporating NHE6 patient variants into the model as altered NHE6 expression, provides a framework for exploring genotype–pH–phenotype correlations. Null mutations (like nonsense, frameshift, splicing, deletions) approximate complete loss of expression, whereas missense mutations can be modelled as partial-to-complete reduction in expression. Conversely, the gain-of-function mutation p.G186R, which is not associated with clinical regression, can be modelled as increased NHE6 expression, resulting in alkaline endosomal pH^45^. Consequently, running the model for a spectrum of expression values of NHE6 (from 0% to 200% with respect to the baseline) shows marked shifts in resultant steady-state endosomal pH values from ∼5.8 to ∼6.8 (*Fig 3D*). Interestingly, these results are consistent with previous literature reporting endosomal pH in non-neuronal cells, which shows that NHE6 overexpression increases luminal pH, whereas NHE6 knockdown induces endosomal hyperacidification across multiple cell types^14,54,55^. This not only supports but also provides independent cross study validation to our modelling results. Additional support for the model derives from endosome experiments in patient-derived neurons harbouring NHE6 mutations, which demonstrate varying degrees of endosomal hyperacidification^56^.

To translate the model to AD, we analysed a publicly available postmortem hippocampal dataset (GSE1297) in which NHE6 expression was downregulated in AD and negatively correlated with Braak staging (*Fig. 3E*). Stratifying individuals by disease severity (control, incipient AD, moderate AD, and severe AD), we observed that estimated endosomal pH was significantly lower—by ∼0.2 pH units—in severe AD compared to controls (*Fig. 3E*). This pH shift is biologically meaningful: even slight alterations can have profound consequences. A notable example is that a 0.2 pH unit perturbation in the cytosol can induce cellular quiescence^57–59^. The model suggests that early-stage NHE6 downregulation may be compensatory, whereas substantial reductions in severe AD may promote maladaptive plasticity, leading to chronic hyperacidification and irreversible progression. We did not observe a significant correlation between age and estimated endosomal pH (*Fig. 3F and Supplementary Fig. 3E*), which may reflect the relatively narrow age range of the postmortem cohort. Notably, when comparing young versus old brain, NHE6 has been reported among the most highly downregulated genes in the old brain^19^, suggesting that age-related NHE6 decline may become more apparent across a broader age spectrum than represented in this dataset. Nevertheless, estimated endosomal pH correlated negatively with postmortem neuropathology, as assessed by neurofibrillary tangle (NFT) score and Braak stage, and positively with antemortem cognitive function as measured by the Mini-Mental State Examination (MMSE) score (*Fig. 3F* and *Supplementary Fig. 3E*).

For further validation, we analysed an independent large postmortem brain dataset (GSE5281) and found that NHE6 expression was significantly downregulated across all brain regions examined, including the entorhinal cortex, hippocampus, medial temporal gyrus, posterior cingulate, superior frontal gyrus, and primary visual cortex, albeit to varying extents (*Fig. 3G*). This regional heterogeneity in NHE6 expression may reflect differences in endosomal function that contribute to region-specific selective vulnerability in AD. The predicted endosomal hyperacidification in AD brains was robustly replicated, with an estimated average decrease in luminal pH of up to ∼0.4 units relative to controls (*Fig. 3G*). These findings support a quantitative link between NHE6 downregulation and endosomal hyperacidification in human AD brains. However, the model likely provides a conservative estimate of endosomal acidification, as pH is regulated by multiple transporters acting in concert, while the model accounts for the NHE6-specific component. For example, previous work reported that APOE4 astrocytes exhibit ∼50% NHE6 reduction—predicting ∼0.26 units of hyperacidification from the model—yet experimentally a decrease of ∼0.84 pH units were observed^20^. This difference likely reflects concurrent NHE9 downregulation observed in APOE4 astrocytes^20^, and the fact that transferrin-based pH measurements capture a mixed pool of early and recycling endosomes^54^. Nevertheless, lentiviral mediated ectopic expression of NHE6 rescued Aβ clearance in APOE4 astrocytes by correcting endosomal pH by ∼0.2 units (of the total ∼0.84-unit shift)^20^, validating that the model, although parameterized with neuronal data, faithfully recapitulates the NHE6-specific contribution.

Overall, the model provides a conceptual framework for defining a reversible therapeutic window and an irreversible maladaptive phase, explaining both the natural history of CS and the potential contribution of NHE6 downregulation to AD pathophysiology. Although our cross-sectional analyses do not directly establish temporal progression or causality, the late-onset regression in CS, the stepwise decline in NHE6 expression and estimated endosomal pH across Braak stages in AD—combined with progressive pathology in NHE6-null mice^25^—collectively support the plausibility of a progressive mechanism. Sustained NHE6 downregulation in AD may contribute to endosomal hyperacidification, potentially leading to maladaptive plasticity and neurodegeneration. Direct longitudinal human studies will be necessary to fully delineate the therapeutic window. Collectively, these findings position endosomal pH as a central node linking NHE6 downregulation to AD, providing a quantitative basis for pH-targeted interventions aimed at preventing maladaptive plasticity before this transition occurs.

### NHE6 Depletion Associates with Aβ Oligomerization and Maladaptive Phenotypic Plasticity

Having established that NHE6 downregulation quantitatively predicts endosomal hyperacidification in AD, we next asked whether this pH shift is mechanistically sufficient to alter Aβ metabolism and trigger maladaptive phenotypic plasticity. We focused on Aβ oligomer (AβO) formation, as metastable AβOs are more potent than monomers or fibrils in inducing synaptic dysfunction and tau pathology^27^. In neurons, the endosomal-lysosomal system serves as a major site for the assembly of pathologically relevant AβOs^27^. Mechanistically, NHE6 downregulation influences Aβ oligomer formation through two pH-sensitive pathways. First, acidic endosomal pH indirectly promotes AβO formation by enhancing Aβ production via modulation of β-secretase (BACE) activity, consistent with its acidic pH optimum, thereby providing more substrate for oligomerization^26^. Second, acidic pH directly promotes AβO formation by accelerating the aggregation of Aβ monomers into oligomers. The endosome is a recognized site for Aβ oligomer nucleation, and acidic pH favors conformational changes that increase Aβ aggregation propensity^27^.

We derived quantitative relationships between pH and each process by fitting experimental data, as detailed in the Methods section. Importantly, BACE activity exhibited a characteristic bell-shaped dependence on pH; specific activity was fitted to a Gaussian distribution with maximal activity at pH 4.5^26^. For Aβ oligomerization, experimental data revealed that the oligomerization rate constant increases exponentially as pH decreases, and this relationship was fitted as linear on a logarithmic scale^27^. Next, using physiological pH values, we simulated the biochemical environment of different cellular compartments where Aβ production and aggregation occur^60^. Two key observations emerged. First, although the pH of recycling endosomes is 0.2 unit higher than that of early endosomes, this shift results in a ∼50% reduction in both BACE activity and AβO formation. Second, at lysosomal pH (4.7), both AβO formation and BACE activity are several orders of magnitude faster than at interstitial pH (7.4) (*Fig. 4A* and *Supplementary Fig. 3F*).

**Figure 4:**
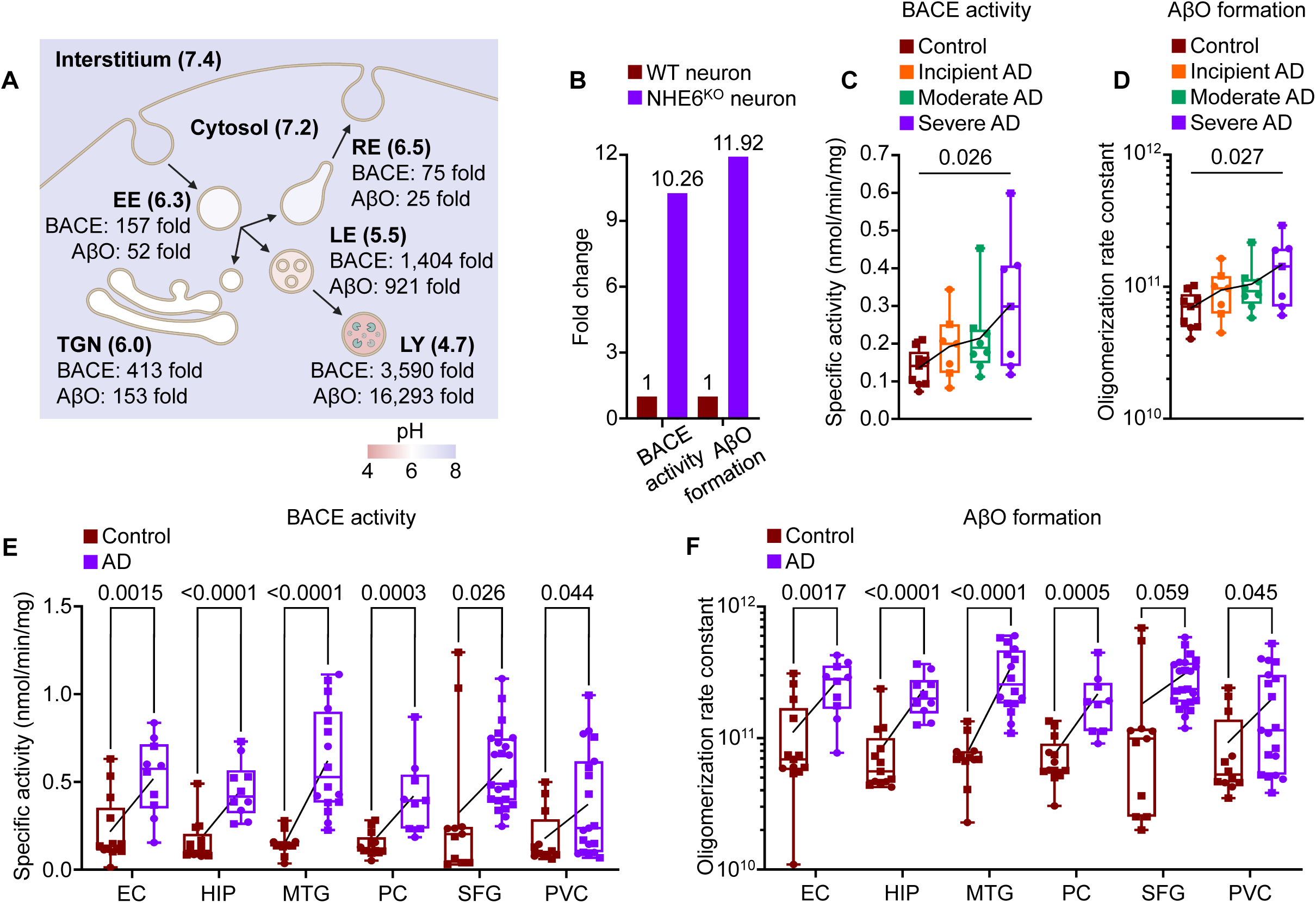
Endosomal hyperacidification promotes Aβ oligomer formation through dual pH-sensitive mechanisms. (A) Schematic showing pH values of different organelles and compartments from the literature^60^ and calculated resultant effects on BACE activity and Aβ oligomer formation. Fold changes relative to interstitial pH (7.4) are shown. Note that recycling endosomes (RE), early endosomes (EE), trans-Golgi network (TGN), late endosomes (LE), and lysosomes (LY) show progressively increasing amyloidogenic activity. The illustration was created with BioRender. (B) Bar plot showing BACE activity and AβO formation in NHE6 knockout neurons (pH 5.88) relative to wild-type (pH 6.57), demonstrating >10-fold increases in both parameters resulting from endosomal hyperacidification. (C, D) Box-whisker plots showing analysis of hippocampal brain dataset GSE1297, where specific BACE activity (C) and Aβ oligomerization rate constant (D) were calculated from endosomal pH derived from NHE6 expression levels. Both measures increase progressively with disease severity (severe AD > moderate AD > incipient AD > control). P values were calculated by one-way ANOVA. (E, F) Box-whisker plots showing validation in independent dataset GSE5281, demonstrating NHE6 downregulation-associated elevated BACE activity (E) and Aβ oligomerization rate constant (F) in AD brains relative to controls across six brain regions: entorhinal cortex (EC), hippocampus (HIP), medial temporal gyrus (MTG), posterior cingulate (PC), superior frontal gyrus (SFG), and primary visual cortex (PVC). P values were calculated by unpaired, two tailed t test. Note that NHE6 downregulation influences Aβ oligomer formation in AD through two pH-dependent mechanisms: increased BACE activity (enhancing substrate availability) and increased Aβ oligomerization rate (accelerating aggregation kinetics). Each scatter point in (C-F) represents an individual postmortem brain; squares represent males and circles represent females, and a line connecting the average values is overlaid. See also Supplementary Figure 3.

We next translated these predictions to NHE6 knockout neurons (wild-type pH 6.57; KO pH 5.88)^61^ and found substantial increases in amyloidogenic activity: a 10.26-fold increase in BACE activity and an 11.92-fold increase in AβO formation (*Fig. 4B*). These results suggest that NHE6 loss alone may promote both elevated Aβ production and rapid oligomerization within the endosomal lumen, independent of changes in APP expression. We then applied our endosome-calibrated model to a postmortem brain dataset containing control and AD brains of varying severity (GSE1297). For each individual, endosomal pH was estimated from NHE6 expression using our model, and specific BACE activity and AβO oligomerization rate were calculated. Both measures increased with disease severity (severe AD > moderate AD > incipient AD > control) (*Fig. 4C-D*). Analysis of an independent postmortem brain dataset validated that pH-regulated BACE activity, and AβO levels are elevated in AD brains across different regions (*Fig. 4E-F*).

To determine whether NHE6-associated Aβ oligomerization correlates with maladaptive phenotypic plasticity in human AD brains, we examined calcium signalling, a major downstream effector of AβO toxicity^62^. Notably, endosomal pH alterations have been shown to affect calcium signalling and phenotypic plasticity in cancer cells^37,38^. Endosomal pH modulation by NHE6 and NHE9 may regulate cytosolic Ca² levels through pathways that include regulation of the plasma membrane expression of Ca² channels, although the precise molecular mechanism(s) remain unclear^63,64^. However, regardless of the direct effects of NHE6 on Ca² homeostasis, calcium signalling serves as a recognised mediator for AβO-mediated neuronal dysfunction and may act both downstream of disease severity and as a contributor to neuronal loss^62^. Accordingly, we evaluated correlations between predicted AβO rate constants and the activity (measured by ssGSEA; see Methods) of two Ca² signalling gene sets, defined based on their up- or down-regulation patterns in AD, as a proxy for maladaptive phenotypic plasticity in AD brains. Given the potential role of NHE6 in synaptic function^61,65^, we extended our analysis to two synapse-related gene sets, also defined by their differential expression in AD. Of note, NHE6 is not part of the *a priori* curated calcium signalling or synapse-related gene sets used in our ssGSEA analysis. As a starting point, to assess whether the coordinated downregulation of Ca² signalling- and synapse-related gene sets in AD simply reflects neuronal loss, we examined their normal expression patterns using the BrainSpan atlas. Most genes showed coordinated developmental profiles, with NHE6 clustering with them. Notably, NHE6 showed strong correlations with genes encoding metabotropic glutamate receptor 5 (*GRM5*; Ca² signaling) and neurexin 1 (*NRXN1*; synapse-related) across both prenatal and postnatal development (*Supplementary Fig. 4A-D*). This co-expression in normal human brain supports a functional link between NHE6 and these pathways, independent of neuronal attrition, and justifies examining NHE6-associated AβO estimates in relation to activity of these gene sets in AD.

In the hippocampal dataset (GSE1297), ssGSEA scores for downregulated gene sets—both Ca² signaling and synapse-related—correlated negatively with neuropathological severity (NFT score, Braak stage) and positively with cognitive function (MMSE score), whereas upregulated gene sets showed the opposite pattern (*Fig. 5A*). Importantly, the NHE6-associated AβO rate constant showed a positive association with ssGSEA scores for upregulated Ca² signalling gene set (R = 0.772, P = 3.6 × 10 ) and a negative association with downregulated Ca² signalling gene set (R = −0.908, P = 1.9 × 10 ¹²) (*Fig. 5B*). Brains with higher inferred AβO burden are likely to show upregulation of stress-associated Ca² genes (e.g., *ITPR2*, *ATP2A3*, *NFATC1*) and downregulation of protective/homeostatic Ca² genes (e.g., *ATP2B2*, *CALM1*, *PPP3R1*). This expression profile, indicative of Ca² signalling remodelling, is reminiscent of maladaptive plasticity^66^. In contrast, the AβO rate constant showed no significant association with upregulated synapse-related gene set (R = −0.034, P = 0.86) but was negatively associated with downregulated synapse-related gene set (R = −0.878, P = 9.0 × 10 ¹¹) (*Fig. 5C*). To validate these findings, we analysed an independent AD and control dataset (GSE5281). The AβO rate constant was positively associated with ssGSEA scores for upregulated Ca² signalling gene set across all examined brain regions, with Pearson correlation coefficients reaching as high as 0.949 (*Supplementary Fig. 5A*). By contrast, associations with upregulated synapse-related gene set were either positive (in middle temporal gyrus and posterior cingulate cortex) or not significant (in all other regions) (*Supplementary Fig. 5B*). For downregulated gene sets—both Ca² signalling and synapse-related— associations were either non-significant (in hippocampus) or negative (in all other regions) (*Supplementary Fig. 5C-D*). Collectively, these results support the plausibility that AβO-associated dysregulation of Ca² signalling and synaptic dysfunction is regionally heterogeneous.

**Figure 5:**
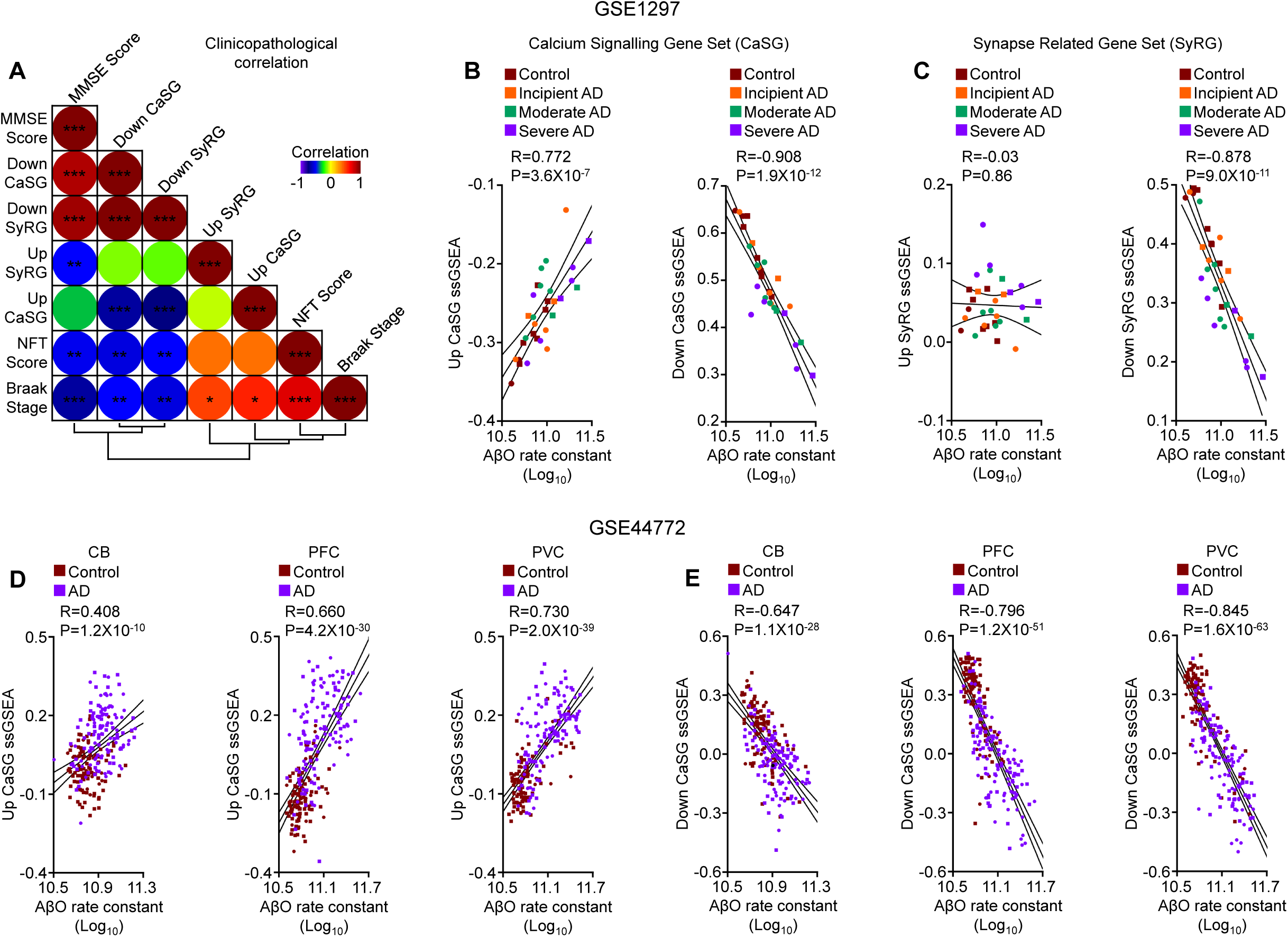
NHE6-associated Aβ oligomerization correlates with dysregulation of calcium signaling and synaptic dysfunction in Alzheimer’s disease. (A) Heatmap of the correlation coefficient matrix depicting clinicopathological associations of ssGSEA scores for upregulated and downregulated Ca² signalling gene sets (Up CaSG and Down CaSG) and upregulated and downregulated synapse-related gene sets (Up SyRG and Down SyRG) in AD. Note that ssGSEA scores for both Down CaSG and Down SyRG correlated positively with antemortem cognitive function (MMSE score) and negatively with postmortem neuropathology (NFT score and Braak stage), whereas upregulated gene sets showed the reciprocal pattern, albeit to varying degrees. *P < 0.05, **P < 0.01, ***P < 0.001 for each comparison. (B) Scatter plots showing that the NHE6-associated AβO rate constant is positively correlated with ssGSEA scores for the upregulated calcium signalling gene set (*left*) and negatively correlated with the downregulated calcium signalling gene set (*right*). Linear fit, Pearson correlation (R), and 95% confidence interval band are shown. (C) Scatter plots showing that the NHE6-associated AβO rate constant had no significant correlation with ssGSEA scores for the upregulated synapse-related gene set (*left*) but was negatively correlated with the downregulated synapse-related gene set (*right*). Linear fit, Pearson correlation (R), and 95% confidence interval band are shown. Data in (A-C) are from the hippocampal brain dataset GSE1297. (D-E) Scatter plots showing independent validation in the GSE44772 dataset, with correlations between the NHE6-associated AβO rate constant and ssGSEA scores for the (D) upregulated and (E) downregulated Ca² signalling gene sets across three brain regions: cerebellum (CB), dorsolateral prefrontal cortex (PFC), and primary visual cortex (PVC). Linear fit, Pearson correlation (R), and 95% confidence interval band are shown. Positive correlations were observed for the upregulated Ca² signalling gene set and negative correlations for the downregulated Ca² signalling gene set across all brain regions, although less pronounced in the cerebellum, consistent with maladaptive calcium-signalling remodelling exhibiting regional heterogeneity. Each scatter point in (D-E) represents an individual postmortem brain; squares represent males and circles represent females. See also Supplementary Figures 4, 5, and 6.

To further explore regional heterogeneity in endosomal dysfunction, we extended our analysis to a large control and AD dataset (GSE44772). The cerebellum—a region relatively resistant to late-onset AD^67^— showed the least pronounced NHE6 downregulation and predicted endosomal hyperacidification compared with the dorsolateral prefrontal cortex and primary visual cortex (*Supplementary Fig. 6A*). These regional differences were also reflected in estimated BACE activity and AβO rate constants (*Supplementary Fig. 6B*). Furthermore, correlations between NHE6-associated AβO rate constants and ssGSEA scores for calcium signalling and synapse-related gene sets were largely consistent with our earlier analysis, except for a significant negative association with upregulated synapse gene set (*Fig. 5D-E and Supplementary Fig. 5C-D*). This contrasts with the non-significant or positive associations observed in other datasets (*Fig. 5C and Supplementary Fig. 6B*), suggesting that AβO-associated synaptic gene remodelling may be context-dependent. Collectively, these findings provide quantitative evidence that NHE6 downregulation-associated endosomal hyperacidification in AD may be sufficient to promote pathological increases in both Aβ production and oligomerization, as well as gene expression changes indicative of maladaptive plasticity.

Experimental evidence supports model predictions. First, NHE6 protein levels were significantly lower in postmortem AD brains than in controls, consistent with the down regulation of its transcript levels in AD observed in microarray analyses^14^. Second, direct measurements of endosomal pH in APOE4 astrocyte and patient fibroblast models of AD showed significant hyperacidification of endosomal pH associated with NHE6 downregulation^20^. Third, NHE6 knockout mice showed increased brain Aβ40 levels and diminished brain weight, suggesting an underlying neurodegenerative pathology^20^. Fourth, and importantly, NHE6 knockdown (∼70%) in a cell culture model of AD led to a ∼1.7-fold increase in Aβ, whereas overexpression reduced it, placing endosomal pH upstream of amyloid pathology^14^. Beyond the studies informing our model, there is further evidence to support acidic pH-dependent increases in BACE activity and Aβ aggregation^68,69^. This is corroborated by in vivo findings from an NHE6-null rat model, which demonstrated age-dependent Aβ aggregation in the hippocampus and cortex^30^. Endosomal hyperacidification-mediated Aβ oligomer formation could in turn induce tau missorting, linking NHE6 loss to tau pathology reported in patient brains as well as rat and human neurons^27,30,48,70^.

The opposing effects of NHE6 depletion (endosomal hyperacidification) and overexpression (endosomal alkalinization) on Aβ levels can be explained, at least in part, by the pH dependence of BACE activity^14^. However, direct experimental evidence linking NHE6 to BACE activity and Aβ oligomerization in neurons remains an important area for future investigation. In summary, the nonlinear relationship between NHE6 levels and endosomal pH exhibits threshold-like behavior. Because Aβ oligomerization is highly pH-sensitive, modest NHE6 reductions may initially trigger compensatory responses. Sustained hyperacidification, however, may promote pathological Aβ aggregation and endosomal dysfunction, pushing neurons into maladaptive states that promote neurodegeneration. Together, these findings establish endosomal pH as a quantitative trigger for Aβ oligomerization and maladaptive phenotypic plasticity in Alzheimer’s disease.

## Discussion

An important conceptual advance of our work is the recognition of endosomal pH as a conserved homeostatic regulator of phenotypic plasticity across two seemingly opposing disease states. In AD, downregulation of NHE6 leads to endosomal acidification, which may initially confer neuroprotection but, upon maladaptation, promotes Aβ oligomerization and neurodegeneration. Conversely, in colorectal cancer, previous work showed that upregulation of the related NHE9 results in endosomal alkalinization, triggering a starvation/pseudo-starvation response that supports cell survival but, when maladaptive, promotes epithelial–mesenchymal transition (EMT) and metastasis^37,38^ (*Fig. 6*). In both contexts, cells respond to stress by shifting their endosomal pH set point; however, this adaptive response ultimately gives rise to more pathological outcomes. This duality lies at the core of the maladaptive plasticity framework and highlights endosomal pH as a unifying mechanistic axis across diverse diseases. Accordingly, therapeutic strategies aimed at restoring physiological endosomal pH may have broad translational potential. Beyond these insights, our findings establish a quantitative framework that integrates genetic, transcriptomic, mathematical, biophysical, and experimental evidence to advance understanding of neurodegenerative processes. Our study has some limitations. First, findings are derived from computational modelling and analyses of public datasets. Second, the pH dependence of BACE activity and Aβ oligomerization is extrapolated from in vitro measurements, which may not fully capture the in vivo environment. Third, direct electrophysiological measurements of NHE6 activity are currently lacking. The model could be refined by incorporating such measurements, along with the contributions of Ca² dynamics and other ion transporters and channels involved in endosomal homeostasis. Despite these limitations, several important implications emerge.

**Figure 6:**
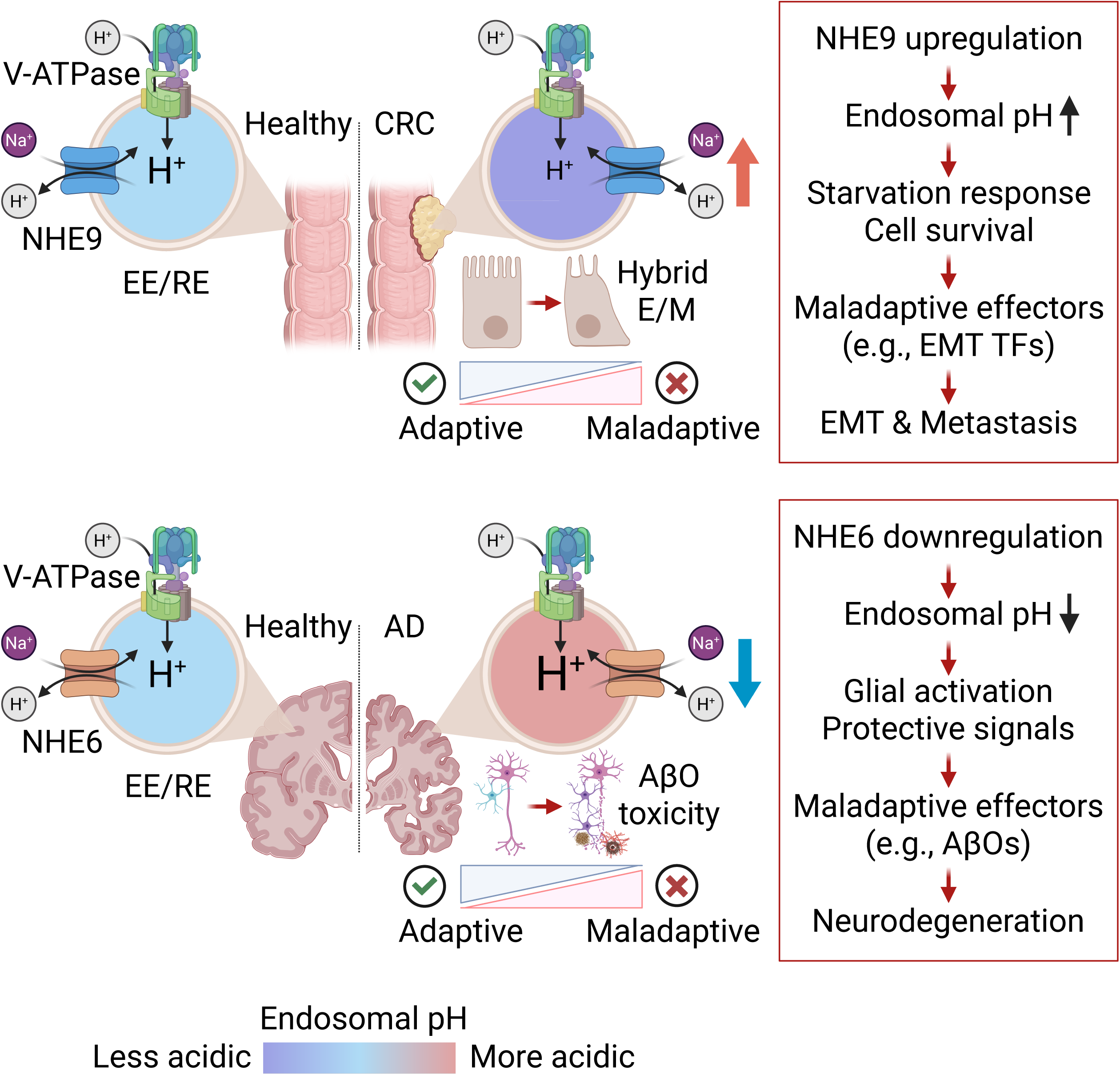
Endosomal pH as a conserved homeostat governing adaptive-to-maladaptive transitions in neurodegeneration and cancer. Schematic illustrating the maladaptive phenotypic plasticity framework across two disease contexts. Top (colorectal cancer (CRC)/malignancy): In cancer cells, NHE9 upregulation promotes endosomal alkalinization, promoting survival through a starvation/pseudo-starvation response. Sustained elevation of endosomal pH induces gene expression programs that modulate cellular plasticity through maladaptive effectors, including epithelial–mesenchymal transition transcription factors (EMT TFs), which underpin hybrid epithelial/mesenchymal (E/M) phenotypes and metastatic progression. Bottom (Alzheimer’s disease (AD)/neurodegeneration): In the brain, NHE6 downregulation—driven by APOE4, aging, or other stressors—causes endosomal hyperacidification. This response may initially be adaptive by enhancing glial activation and neuroprotective signals, including beneficial endosomal function and cargo clearance. However, progressive hyperacidification acts through maladaptive effectors such as Aβ oligomers (AβOs), through both increased BACE activity and direct pH-dependent acceleration of oligomerization kinetics, which underpin AβO toxicity and neurodegeneration. This leads to a pathogenic model predicting that, in both disease contexts, cells respond to stress by recalibrating their endosomal pH set point, but this initial adaptive response may ultimately become maladaptive through mechanisms including calcium dyshomeostasis and synaptic dysfunction. The threshold-like behavior may define a therapeutic window for interventions correcting endosomal pH before irreversible pathology develops. The illustration was created with BioRender.

First, a notable example of the adaptive-to-maladaptive trajectory in AD involves NHE6-regulated recycling of low-density lipoprotein receptor-related protein 1 (LRP1)^20^. Under physiological conditions, recycling endosomes are ∼0.2 pH units more alkaline than early endosomes, whereas late endosomes are ∼0.8 pH units more acidic^60^ (*Fig. 4A*). This observation is not merely descriptive but functionally instructive, leading to a luminal pH hypothesis of endocytic recycling: modest alkalization favors recycling, whereas hyperacidification promotes degradative sorting. Consistent with this, it was shown that NHE6 downregulation and resulting endosomal hyperacidification inhibit LRP1 recycling in APOE4 astrocytes, reducing its surface expression^20^. Because LRP1 is a receptor for both APOE and Aβ, this reduction may have biphasic effects: initially mitigating APOE4-induced cholesterol and trafficking deficits (beneficial), but later impairing Aβ clearance as disease progresses (detrimental)^20,71^. Importantly, NHE6 restoration rescued surface LRP1 and improved Aβ clearance, supporting the reversibility of this maladaptive transition^20^.

Second, building on the ideas outlined above, we propose that endosomal dysfunction in AD may broadly follow an adaptive-to-maladaptive progression. Early in the disease, presymptomatic amplification of endosomes in both size and number, along with increased endocytosis^3^, could represent a remodeling mechanism aimed at enhancing cargo clearance. However, as pathology progresses and endosomal function falls below a critical tipping point, this response may become maladaptive. As a result, the endosome transitions from a dynamic sorting hub to a trafficking bottleneck, with endosomal dysfunction emerging as an active promoter of disease progression. The existence of such a threshold in our pathogenic model suggests a potential therapeutic window before maladaptation becomes entrenched. This framework also helps explain the clinical heterogeneity observed in AD and CS: variable rates of NHE6 loss or compensatory responses could shift the timing of the maladaptive transition. Restoring physiological endosomal pH, therefore, represents a promising and broadly applicable therapeutic strategy.

Third, the reciprocal enrichment pattern of NHE6 in neurons and NHE9 in glia across multiple human brain datasets highlights a cellular basis for endosomal pH regulation in the central nervous system. This suggests that neurons and glia may have evolved distinct molecular mechanisms to regulate endosomal acidity, likely reflecting their unique physiological demands. Nevertheless, this specialization is not absolute: NHE6 has been shown to play a role in astrocytes^20^, and NHE9 has also been reported to function in neurons^11^, indicating functional overlap that may contribute to context-dependent cellular responses. The enrichment of SNARE signaling, synaptogenesis, and glutamatergic receptor signaling among NHE6-coexpressed genes further underscores its neuronal functional context. Extending beyond AD pathology, NHE6 also exhibits a co-expression pattern with Ca² signalling genes in the Developing Human Brain Atlas and the Aging, Dementia, and TBI Study datasets, providing independent support for the importance of endosomal pH in Ca² signalling in human (patho)physiology (*Supplementary Fig. 4A-B and Supplementary Fig. 7A*). Accordingly, loss of NHE6 expression may represent a potential determinant of neuronal vulnerability in AD pathogenesis.

Fourth, systematic curation of 120 NHE6 patient variants reveals that null mutations invariably cause severe phenotypes and are associated with clinical regression, whereas the only gain-of-function mutation (p.G186R) lacks regression, indicating that loss of function promotes endosomal hyperacidification, maladaptive plasticity, and neurodegeneration distinct from neurodevelopmental defects^12,45^. Although no NHE6 variants have yet been linked to AD, large-scale sequencing efforts will likely identify both common and rare variants, increasing the risk of misleading pathogenicity assignments. For example, beyond our curated dataset, additional missense variants of uncertain significance in NHE6 have been reported in epilepsy patients (hemizygous: p.R3L, p.R3W, p.L90P, p.F151L; heterozygous: p.H121R, p.Y162C, p.I238V, p.V316M, p.S615G)^72^. Functional validation of patient variants is therefore imperative, as illustrated by the p.A9S variant in NHE6—present in a male with Angelman-like syndrome but absent in his affected sister, strongly suggesting an unrelated etiology—which proved functionally neutral in both cellular and mouse models^41,73,74^.

Fifth, our mathematical model of neuronal endosomal pH regulation, applied to postmortem brain data, represents a significant advance by integrating major ion fluxes into a predictive framework. The extreme pH sensitivity of Aβ oligomer formation—which increases by several orders of magnitude from extracellular to lysosomal pH—together with the threshold-like behavior of NHE6 expression defines a quantifiable region of sensitivity for disease progression. This raises a question: is acidic pH-mediated aggregation universal to all amyloids? Notably, human islet amyloid polypeptide (hIAPP)—which links type 2 diabetes to AD—behaves differently: acidic pH inhibits its aggregation due to a conserved histidine in its core (H18; ConSurf score 7) whose protonation introduces inter-chain repulsion^75^. In contrast, Aβ contains three low-to-moderately conserved histidines in its N-terminus (H6, H13, and H14; ConSurf scores 3, 2, and 4), suggesting a regulatory rather than a fundamental role in oligomer formation: their protonation relieves electrostatic repulsion, allowing the conserved hydrophobic C-terminus to promote oligomerization^69^ (*Supplementary Fig. 7B-D*). Thus, Aβ oligomer formation emerges as a key mechanistic link between endosomal hyperacidification and AD progression.

In conclusion, this work reframes the role of NHE6 in AD from a simple linear pathway to a dynamic, biphasic promoter of maladaptive phenotypic plasticity. By integrating a quantitative mathematical model with validation and evolutionary parallels from cancer biology, we identify endosomal pH as a systems-level regulator governing pathological cell-state transitions. This model positions NHE6 as an upstream druggable node driving amyloid-mediated neurodegeneration, influencing Aβ oligomer formation through both increased substrate availability and accelerated aggregation kinetics. Critically, even a pH shift of 0.2 units is predicted to produce a twofold change in Aβ oligomer formation. Our findings are hypothesis-generating and provide a foundation for future studies, with significant therapeutic implications for defining a critical window for intervention in both CS and AD. While reversing neurodevelopmental defects that arise during early brain development may be challenging, early detection and correction of endosomal pH defects could offer a therapeutic opportunity to mitigate the progressive neurodegeneration that characterizes late-stage CS. In the context of AD, the adaptive-to-maladaptive transition is expected to occur insidiously over time, rather than abruptly, as NHE6 expression progressively declines. This highlights that early strategies aimed at stabilizing endosomal pH could prevent progression to an irreversible pathological state. Endosomal pH emerges as an evolutionarily conserved regulator of maladaptive phenotypic plasticity across both cancer and neurodegeneration, offering a unifying framework for understanding how cellular adaptations to stress can ultimately promote disease progression.

## Methods

### 1. Mathematical modelling of NHE6-regulated endosomal pH homeostasis

We used a computational framework to examine how NHE6 abundance influences neuronal endosomal pH regulation. The model builds upon the mathematical framework originally developed by Ishida et al. for lysosomal acidification^53^, and subsequently extended to describe NHE9-dependent endosomal pH regulation in cancer cells^37^. In the present study, we adapted this framework to incorporate the physiological context of NHE6 in neuronal endosomes. The model incorporates the primary ion transport mechanisms governing endosomal acidification, including V-ATPase-mediated proton pumping, NHE6-mediated proton leak, CLC Cl /H antiport, passive ion leakage across the endosomal membrane, and other ion regulating elements^53^. Each transport process is represented as an independent module, and these modules are combined into a coupled system of differential equations describing the net flux of each ionic species into and out of the endosomal lumen. The model was solved numerically to obtain steady-state luminal pH as a function of NHE6 expression level. For all simulations, the endosomal system is assumed to reach steady state at 3000 seconds, which is consistent with the time taken for transferrin uptake to reach steady state (∼1 hour)^76^ and to show colocalization with NHE6^14^. The marginal gain in pH per 50% increase in NHE6 (ΔpH/ΔNHE6) was calculated as the difference in steady-state endosomal pH between sequential increases in NHE6 expression. Model parameter values are provided in *Supplementary Table 2*.

#### V-ATPase mediated proton pump

The V-ATPase activity was represented by a single-pump proton transport flux , with positive values membrane potential (ΔΨ) was parameterized using published current–voltage measurements^77^. To compute the total V-ATPase-mediated proton flux, the single-pump rate is scaled by , which denotes the number of active V-ATPase complexes on the endosomal membrane.

#### NHE6 mediated proton leak

NHE6 was modelled as an electroneutral Na /H exchanger that provides a counterbalancing proton efflux pathway opposing V-ATPase-mediated acidification. Although NHE6 can mediate the exchange of luminal H with either cytosolic Na or K , in this model it was represented specifically as coupling Na influx to H efflux from the lumen. The exchanger activity was assumed to increase under more acidic luminal conditions, thereby limiting excessive proton accumulation and stabilizing endosomal pH. The proton flux mediated by NHE6 is described by the following equation:

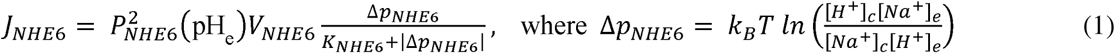

The NHE model developed by Marcoline et al. for the plasma membrane NHE homolog was adapted for NHE6^78^. We modified the original model, and instead of a linear drop from 1 to 0 over the pH range of 6.55–7.25, the term _NHE6_ pH was modeled using the following sigmoidal function.

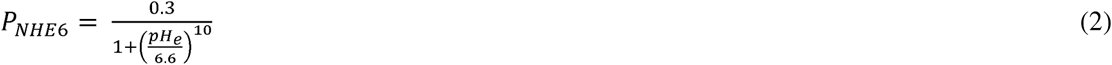

A maximum turnover rate of 1500 s ¹ was assigned to _NHE6_, based on reported values for the bacterial Na /H exchanger NhaA^79,80^. Although members of the cation/proton antiporter superfamily share low sequence similarity, they exhibit a conserved three-dimensional fold, and the two-domain elevator mechanism of transport is maintained from bacteria to mammals, supporting the use of bacterial estimates for NHE6^79,81^. This high turnover rate explains why NHE6 can overwhelm the capacity of the proton pump V-ATPase, such that even small alterations in the NHE6 leak pathway can result in significant pH shifts. *K*_NHE6_ was set to 3 *k_B_T* as previously described^78^. To determine the total flux, the activity of a single exchanger was multiplied by the total number of exchangers.

#### CLC Cl^-^/H^+^ antiporter

The CLC component was modelled using the 2:1 Cl :H exchange stoichiometry reported for CLC7-like transporters. Using electrophysiological recordings of CLC7, Ishida et al.^53^ derived an empirical expression for single-transporter flux that depends on both the pH gradient and the Cl concentration ratio across the endosomal membrane. This expression takes the following form:

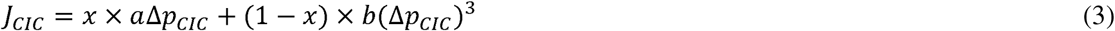

Here, *a* is 0.3, *b* is 1.5 10^−5^, Δ*p_cic_* is the driving force defined as:

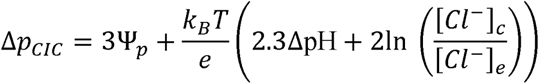

and, *x* is a switching function defined as:

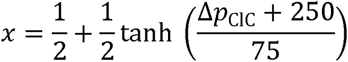

To compute the total fluxes, the activity of a single CLC antiporter is multiplied by both the total number of antiporters (*N*_ClC_) and the respective stoichiometric coefficients—2 for Cl influx and 1 for H efflux.

#### Membrane potential

Following the framework established by Ryback et al.^82^, the membrane potential from the net charge density within the endosome was calculated using the following equation:

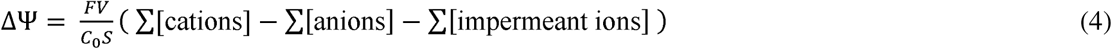

In this equation, is Faraday’s constant, is the membrane capacitance per unit area, and and are endosomal surface area and volume, respectively. The summation terms represent the total concentrations of permeable cations and anions, while impermeant ions (Donnan particles, “B”) accounts for negatively charged luminal macromolecules.

#### Passive leaks for H^+^, Na^+^, K^+^, and Cl^-^ ions

Passive leaks were modelled using the following general expression^46^:

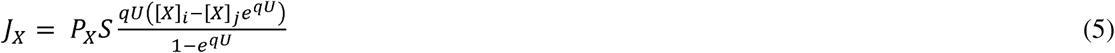

In this expression, *q* denotes the ionic charge, *S* is membrane surface area, U = eψ/*k_g_T* is the reduced membrane potential, and [*X*] is the concentration of ion *X* across compartments *i* and *j*.

#### Numerical solutions

The temporal evolution of luminal ion concentrations is governed by the following system of differential equations:

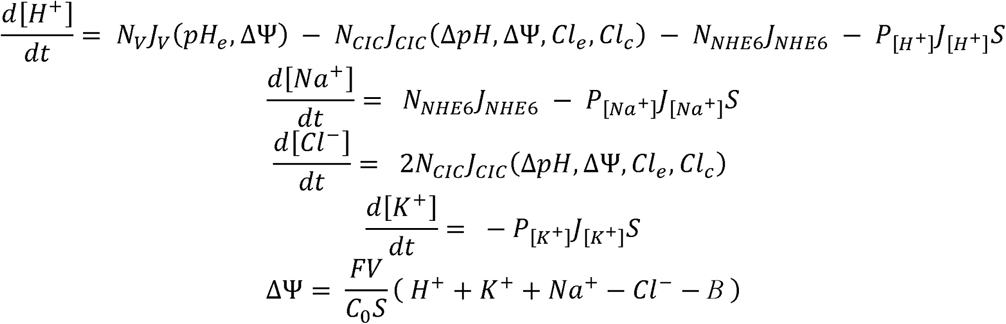

### 2. Mathematical modeling of pH-dependent kinetics

To quantitatively describe the pH dependence of key processes in amyloid pathology, mathematical functions based on empirical data were derived. The catalytic activity of BACE and the oligomerization kinetics of Aβ were each modeled as continuous functions of pH to enable the interpolation at any given pH within the physiological range.

#### pH dependence of BACE catalytic activity

The specific activity of BACE, denoted as (in nmol/min/mg), was modeled as a function of pH using a Gaussian distribution to represent the characteristic bell-shaped dependence of enzyme activity on pH. This model was parameterized using experimental activity data from Vassar et al.^26^, which identified a distinct activity optimum in the acidic range. The fitted curve exhibited a peak amplitude of 9.5 nmol/min/mg at an optimal pH of 4.5 with a standard deviation of 0.715. This Gaussian relationship captures the decline in enzymatic activity observed under both more acidic and more neutral conditions relative to the optimum, described by:

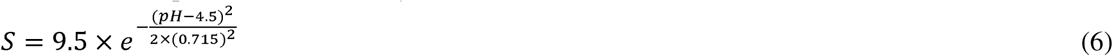

or equivalently,

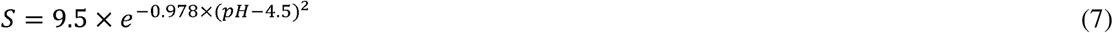

The fold-change in BACE activity for a pH change from pH to pH is derived from the ratio *S* (pH_2_) /*S* (pH_1_). yielding:

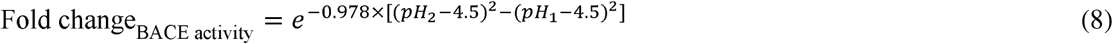

This relationship reflects the relative change in catalytic efficiency as pH deviates from the optimal value of 4.5.

#### pH dependence of Aβ oligomerization kinetics

The Aβ oligomer (AβO) formation rate constant (*K*) was derived from the experimental data of Schützmann et al.^27^ and modeled as a linear function of pH on a logarithmic scale, yielding a slope of −1.56 and an intercept of 21.11:

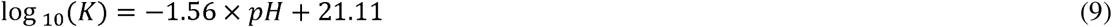

or equivalently,

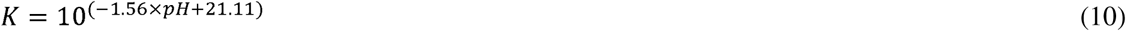

The fold-change in the oligomerization rate for a pH change from *pH* to *pH* is derived from the ratio *K* (pH_2_) /*K* (pH_1_), which simplifies to:

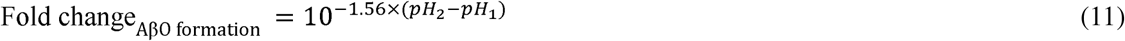

This relationship indicates that one-unit decrease in pH corresponds to an approximately 36-fold increase in the oligomerization rate.

Fold changes for both BACE activity and AβO formation were calculated using the above equations relative to interstitial pH (7.4) for each organellar pH value: recycling endosomes (6.5), early endosomes (6.3), trans-Golgi network (6.0), late endosomes (5.5), and lysosomes (4.7), based on pH values from the published literature^60^. For NHE6 knockout neurons (wild-type pH 6.57, knockout pH 5.88)^61^, fold changes in BACE activity and AβO formation were calculated relative to the wild-type using these equations.

### 3. Single-cell transcriptomic analysis

Single-cell RNA sequencing data were obtained from two independent normal human brain datasets: LIBD (n=4) and IsoHuB (n=10). For cross-disease validation, six additional single-cell datasets from the PsychENCODE Consortium were analyzed, including samples from patients with post-traumatic stress disorder (PTSD), major depressive disorder (MDD), autism spectrum disorder (ASD), Williams syndrome (WS), schizophrenia (SCZ), and bipolar disorder (BD), as well as matched controls: PTSDBrainomics (PTSD n=6, MDD n=4, control n=9), DevBrain-snRNAseq (ASD n=9, WS n=3, control n=4), MultiomeBrain-DLPFC (SCZ n=6, BD n=10, control n=5), UCLA-ASD (ASD n=27, control n=25), CMC (SCZ n=47, control n=53), and SZBDMulti-Seq (SCZ n=24, BD n=24, control n=24). All samples were from adult postmortem dorsolateral prefrontal cortex (DLPFC). UMAP plots and dot plots (mean expression, percent expressing cells, scaled per gene) were extracted for *SLC9A6* and *SLC9A9* from the PsychSCREEN portal. Bar graphs were generated to display mean expression and percentage of expressing cells across annotated cell types.

### 4. Co-expression and Ingenuity Pathway Analysis

Gene–gene co-expression matrices were derived from two independent RNA-seq datasets: the Developing Human Brain Atlas (524 brain samples from 8 postconception weeks to 40 years of age) and the Aging, Dementia, and TBI Study (377 brain samples from aged donors with traumatic brain injury and matched controls). Pearson correlation coefficients were obtained for NHE6 against all other transcripts (52,375 transcripts in the Developing Human Brain Atlas; 50,280 transcripts in the Aging, Dementia, and TBI Study). The top 500 positively correlated genes were selected since they represent ∼1% of human brain transcriptome transcripts for functional enrichment analysis. Ingenuity Pathway Analysis (IPA, Qiagen) was performed on the top ∼500 co-expressed genes, including ties in correlation coefficients, to identify enriched canonical pathways, upstream regulators, and disease associations. Canonical pathway enrichment was assessed using activation z-scores and P value calculated based on the direction of gene expression changes. Upstream regulator analysis was performed using the IPA knowledge base, with activation z-scores indicating predicted activation or inhibition. For disease association, the Analysis Match function in IPA was used to compare upstream regulator activation states between the NHE6 co-expression signature and curated disease-associated gene expression datasets.

### 5. Curation of NHE6 patient variants

A systematic literature search was conducted to identify published NHE6 (*SLC9A6*) patient variants. PubMed and Google Scholar were searched, along with the HGMD, ClinVar, and dbSNP databases. Inclusion criteria were: (1) original research articles reporting human NHE6 variants, (2) clinical phenotype information (when available), and (3) molecular characterization (when performed). Exclusion criteria were: (1) variants reported solely in ClinVar or dbSNP database without an accompanying original research article, (2) variants lacking clearly reported nucleotide or amino acid coordinates. To enable cross-study comparison, all variant coordinates were mapped to a common reference sequence (NM_006359) independent of the transcript identifier used in the original publication; two identified variants specific to the longer NHE6 isoform were mapped to NM_001042537 instead. Data extracted included mutation type, nucleotide change, amino acid change, affected domain, functional consequence (loss-of-function, neutral, or gain-of-function when available), ClinVar/dbSNP entry, and associated clinical phenotypes. The MutationTaster algorithm was used for functional prediction of genetic variants. Transmembrane domain residue locations were assigned based on sequence alignment with the structure of the related endosomal NHE9 (PDB: 8PXB). A total of 120 unique patient variants were curated and compiled into Supplementary Table 1.

### 6. Structural modeling and conservation analysis

The three-dimensional structure of the NHE6 dimer was modelled using the SWISS-MODEL server based on the related NHE9 structure (PDB: 8PXB). Other structures (hIAPP filament, PDB: 9ULZ; Aβ42 filament, PDB: 7Q4B; Aβ42 monomer, PDB: 1Z0Q; Aβ40 monomer, PDB: 1BA4) were also analysed. Evolutionary conservation scores were calculated using the ConSurf Colab server with default parameters, which employs an empirical Bayesian methodology to estimate conservation scores across homologous sequences^83^. Scores range from 1 (variable) to 9 (highly conserved) and were visualized using a colourblind-friendly green-to-purple colour code. Hydrophobicity analysis was performed using the scale of Kessel and Ben-Tal^84^ and visualized using a blue-to-yellow colour code. Missense and nonsense patient variants were mapped onto the modelled structure and visualized as α-carbon spheres using PyMOL, and also shown using a lollipop representation.

### 7. Application of the model to Alzheimer’s disease

Five independent postmortem human brain datasets were analysed. GSE1297 included 31 samples of hippocampal gene expression from individuals classified into four groups: control (n=9), incipient AD (n=7), moderate AD (n=8), and severe AD (n=7). NHE6 expression levels were extracted, and endosomal pH was estimated using the mathematical model described above. Neuropathological assessments included Braak stage and neurofibrillary tangle (NFT) score. Antemortem cognitive function was assessed by Mini-Mental State Examination (MMSE) score. GSE5281 included 161 samples from six brain regions: entorhinal cortex (control n=13; AD n=10), hippocampus (control n=13; AD n=10), medial temporal gyrus (control n=12; AD n=16), posterior cingulate cortex (control n=13; AD n=9), superior frontal gyrus (control n=11; AD n=23), and primary visual cortex (control n=12; AD n=19). Additional datasets derived from a cohort of 230 postmortem brain donors (control, n = 101; AD, n = 129) from GSE44772. This dataset comprises the following subseries: GSE44768 (cerebellum), GSE44770 (dorsolateral prefrontal cortex), and GSE44771 (primary visual cortex). NHE6 expression levels and estimated endosomal pH were compared between control and AD cases. For all datasets, NHE6-regulated BACE catalytic activity and Aβ oligomerization kinetics were predicted based on the estimated endosomal pH.

### 8. ssGSEA scores

Single-sample Gene Set Enrichment Analysis (ssGSEA) scores were obtained using the GSEApy module. For ssGSEA analysis, an a priori curated set of 32 calcium-signalling genes and 42 synapse-related genes was obtained through analysis of the GSE1297 dataset^85^. The calcium-signalling genes were classified into two gene sets: 18 downregulated in AD and 14 upregulated in AD, defined based on their down- or up-regulation expression patterns. Likewise, the synapse-related genes were divided into two gene sets: 22 downregulated and 20 upregulated in AD. The ssGSEA score represents the relative activity level of a pathway, capturing the coordinated expression of genes within each sample.

## 9. Data analysis

Data analyses were performed using GraphPad Prism 9. The statistical tests used are indicated in the figure legends. For all analyses, p-values were two-tailed, and statistical significance was defined as P < 0.05.

## 10. Data Availability

All datasets analysed in this study are publicly available. The LIBD, IsoHuB, and other PsychENCODE Consortium datasets are accessible through the PsychSCREEN portal. Microarray datasets (GSE1297, GSE5281, and GSE44772, the latter comprising GSE44768, GSE44770, and GSE44771) are available through the Gene Expression Omnibus (GEO) database. The Developing Human Brain Atlas is accessible through https://www.brainspan.org/, and the Aging, Dementia, and TBI Study dataset is available through the https://aging.brain-map.org/ portal hosted by the Allen Institute for Brain Science. The NHE9 structural model used for homology modelling is available in the Protein Data Bank (PDB ID: 8PXB). Additional PDB structures analysed in this study include 9ULZ, 7Q4B, 1Z0Q, and 1BA4. All other data generated or analysed during this study are included in the figures and supplementary information files.

## 11. Code availability

The code for the mathematical model for endosomal pH is available at https://github.com/csbBSSE/Endosomal-pH-triggers-A-beta-oligomerisation-and-maladaptive-phenotypic-plasticity-in-AD

## Acknowledgements

H. P. was supported by the Prime Minister Early Career Research Grant (ANRF/ECRG/2024/002999/LS) awarded by the Anusandhan National Research Foundation (ANRF), Govt. of India, and by funds from the Centre for Brain Research (CBR). M.K.J. was supported by Ramanujan Fellowship (SB/S2/RJN-049/2018) awarded by the Science and Engineering Research Board (SERB), Department of Science and Technology, Govt. of India. M.K.J. was also supported by Param Hansa Philanthropies.

## Author contributions

H.P. and M.K.J. conceived the project. R.K.M, A.S.D., and H.P. performed the experiments and analyzed the data. H.P., M.K.J., A.S.D., R.K.M wrote and reviewed the manuscript. All authors have read and approved the manuscript.

## References

1 Selkoe, D. J. & Hardy, J. The amyloid hypothesis of Alzheimer’s disease at 25 years. EMBO Mol Med 8, 595–608, doi:10.15252/emmm.201606210 (2016).

2 Troncoso, J. C. et al. Neuropathology of preclinical and clinical late-onset Alzheimer’s disease. Ann Neurol 43, 673–676, doi:10.1002/ana.410430519 (1998).

3 Nixon, R. A. Amyloid precursor protein and endosomal-lysosomal dysfunction in Alzheimer’s disease: inseparable partners in a multifactorial disease. FASEB J 31, 2729–2743, doi:10.1096/fj.201700359 (2017).

4 Small, S. A. & Petsko, G. A. Endosomal recycling reconciles the Alzheimer’s disease paradox. Sci Transl Med 12, doi:10.1126/scitranslmed.abb1717 (2020).

5 Donowitz, M., Ming Tse, C. & Fuster, D. SLC9/NHE gene family, a plasma membrane and organellar family of Na(+)/H(+) exchangers. Mol Aspects Med 34, 236–251, doi:10.1016/j.mam.2012.05.001 (2013).

6 Fuster, D. G. & Alexander, R. T. Traditional and emerging roles for the SLC9 Na+/H+ exchangers. Pflugers Arch 466, 61–76, doi:10.1007/s00424-013-1408-8 (2014).

7 Pedersen, S. F. & Counillon, L. The SLC9A-C Mammalian Na(+)/H(+) Exchanger Family: Molecules, Mechanisms, and Physiology. Physiol Rev 99, 2015–2113, doi:10.1152/physrev.00028.2018 (2019).

8 Prasad, H. & Rao, R. Endosomal Acid-Base Homeostasis in Neurodegenerative Diseases. Rev Physiol Biochem Pharmacol 185, 195–231, doi:10.1007/112_2020_25 (2023).

9 Prasad, H. & Rao, R. Linking endo-lysosomal pH, sterol, and trafficking to neurodegenerative disease. FEMS Yeast Res 25, doi:10.1093/femsyr/foaf034 (2025).

10 Kondapalli, K. C., Prasad, H. & Rao, R. An inside job: how endosomal Na(+)/H(+) exchangers link to autism and neurological disease. Front Cell Neurosci 8, 172, doi:10.3389/fncel.2014.00172 (2014).

11 Prasad, H. NHE9 and Endosomal pH: Converging Mechanisms in Neurodevelopmental, Psychiatric and Neurodegenerative Disorders. Eur J Neurosci 62, e70334, doi:10.1111/ejn.70334 (2025).

12 Kavanaugh, B. C. et al. Christianson syndrome across the lifespan: genetic mutations and longitudinal study in children, adolescents, and adults. J Med Genet 61, 1031–1039, doi:10.1136/jmg-2024-109973 (2024).

13 Funk, C. C. et al. Mining the gaps: Deciphering Alzheimer’s biology through AI-driven reconciliation. J Prev Alzheimers Dis 13, 100402, doi:10.1016/j.tjpad.2025.100402 (2025).

14 Prasad, H. & Rao, R. The Na+/H+ exchanger NHE6 modulates endosomal pH to control processing of amyloid precursor protein in a cell culture model of Alzheimer disease. J Biol Chem 290, 5311–5327, doi:10.1074/jbc.M114.602219 (2015).

15 Mayburd, A. & Baranova, A. Knowledge-based compact disease models identify new molecular players contributing to early-stage Alzheimer’s disease. BMC Syst Biol 7, 121, doi:10.1186/1752-0509-7-121 (2013).

16 Webster, J. A. et al. Genetic control of human brain transcript expression in Alzheimer disease. Am J Hum Genet 84, 445–458, doi:10.1016/j.ajhg.2009.03.011 (2009).

17 Kundra, R., Ciryam, P., Morimoto, R. I., Dobson, C. M. & Vendruscolo, M. Protein homeostasis of a metastable subproteome associated with Alzheimer’s disease. Proc Natl Acad Sci U S A 114, E5703–E5711, doi:10.1073/pnas.1618417114 (2017).

18 Park, C., Ha, J. & Park, S. Prediction of Alzheimer’s disease based on deep neural network by integrating gene expression and DNA methylation dataset. Expert Systems with Applications 140, 112873 (2020).

19 Naumova, O. Y. et al. Age-related changes of gene expression in the neocortex: preliminary data on RNA-Seq of the transcriptome in three functionally distinct cortical areas. Dev Psychopathol 24, 1427–1442, doi:10.1017/S0954579412000818 (2012).

20 Prasad, H. & Rao, R. Amyloid clearance defect in ApoE4 astrocytes is reversed by epigenetic correction of endosomal pH. Proc Natl Acad Sci U S A 115, E6640–E6649, doi:10.1073/pnas.1801612115 (2018).

21 Xu, P. T. et al. Differences in apolipoprotein E3/3 and E4/4 allele-specific gene expression in hippocampus in Alzheimer disease. Neurobiol Dis 21, 256–275, doi:10.1016/j.nbd.2005.07.004 (2006).

22 Lee, J. H. et al. Lysosomal proteolysis and autophagy require presenilin 1 and are disrupted by Alzheimer-related PS1 mutations. Cell 141, 1146–1158, doi:10.1016/j.cell.2010.05.008 (2010).

23 Lee, H. et al. ApoE4-dependent lysosomal cholesterol accumulation impairs mitochondrial homeostasis and oxidative phosphorylation in human astrocytes. Cell Rep 42, 113183, doi:10.1016/j.celrep.2023.113183 (2023).

24 Prasad, H. Bridging LysoPD and MitoPD: Lysosomal pH Links Two Hallmarks of Parkinson’s Disease. Mov Disord doi:10.1002/mds.70492 (2026).

25 Stromme, P. et al. X-linked Angelman-like syndrome caused by Slc9a6 knockout in mice exhibits evidence of endosomal-lysosomal dysfunction. Brain 134, 3369–3383, doi:10.1093/brain/awr250 (2011).

26 Vassar, R. et al. Beta-secretase cleavage of Alzheimer’s amyloid precursor protein by the transmembrane aspartic protease BACE. Science 286, 735–741, doi:10.1126/science.286.5440.735 (1999).

27 Schutzmann, M. P. et al. Endo-lysosomal Abeta concentration and pH trigger formation of Abeta oligomers that potently induce Tau missorting. Nat Commun 12, 4634, doi:10.1038/s41467-021-24900-4 (2021).

28 Xian, X. et al. Reversal of ApoE4-induced recycling block as a novel prevention approach for Alzheimer’s disease. Elife 7, doi:10.7554/eLife.40048 (2018).

29 Pohlkamp, T. et al. NHE6 depletion corrects ApoE4-mediated synaptic impairments and reduces amyloid plaque load. Elife 10, doi:10.7554/eLife.72034 (2021).

30 Lee, Y. et al. Early lysosome defects precede neurodegeneration with amyloid-beta and tau aggregation in NHE6-null rat brain. Brain 145, 3187–3202, doi:10.1093/brain/awab467 (2022).

31 Ren, W. et al. Ketamine promotes the amyloidogenic pathway by regulating endosomal pH. Toxicology 471, 153163, doi:10.1016/j.tox.2022.153163 (2022).

32 Huang, N. et al. Targeting the HDAC4-NHE6-endosomal pH axis restores amyloid-beta clearance and cognitive function in Alzheimer’s disease mice. J Nanobiotechnology, doi:10.1186/s12951-026-04297-2 (2026).

33 Kulkarni, P. & Salgia, R. Comprehending phenotypic plasticity in cancer and evolution. iScience 27, 109308, doi:10.1016/j.isci.2024.109308 (2024).

34 Hari, K., Ullanat, V., Balasubramanian, A., Gopalan, A. & Jolly, M. K. Landscape of epithelial-mesenchymal plasticity as an emergent property of coordinated teams in regulatory networks. Elife 11, doi:10.7554/eLife.76535 (2022).

35 Garcia-Jimenez, C. & Goding, C. R. Starvation and Pseudo-Starvation as Drivers of Cancer Metastasis through Translation Reprogramming. Cell Metab 29, 254–267, doi:10.1016/j.cmet.2018.11.018 (2019).

36 Prasanna, C. V. S., Jolly, M. K. & Bhat, R. Dependence of mesenchymally transitioned tumor niche fitness on cell-cell and cell-matrix adhesions. Biophys J, doi:10.1016/j.bpj.2026.01.002 (2026).

37 Prasad, H. et al. An Endosomal Acid-Regulatory Feedback System Rewires Cytosolic cAMP Metabolism and Drives Tumor Progression. Mol Cancer Res 22, 465–481, doi:10.1158/1541-7786.MCR-23-0606 (2024).

38 Prasad, H. et al. Endosomal pH is an evolutionarily conserved driver of phenotypic plasticity in colorectal cancer. NPJ Syst Biol Appl 10, 149, doi:10.1038/s41540-024-00463-0 (2024).

39 Prasad, H. Genes for endosomal pH regulators NHE6 and NHE9 are dysregulated in the substantia nigra in Parkinson’s disease. Gene 927, 148737, doi:10.1016/j.gene.2024.148737 (2024).

40 Prasad, H. & Rao, R. Histone deacetylase-mediated regulation of endolysosomal pH. J Biol Chem 293, 6721–6735, doi:10.1074/jbc.RA118.002025 (2018).

41 Ilie, A. et al. Assorted dysfunctions of endosomal alkali cation/proton exchanger SLC9A6 variants linked to Christianson syndrome. J Biol Chem 295, 7075–7095, doi:10.1074/jbc.RA120.012614 (2020).

42 Jiao, J. P. et al. Missense variants in SLC9A6 cause partial epilepsy without neurodevelopmental delay. Orphanet J Rare Dis 20, 380, doi:10.1186/s13023-025-03924-9 (2025).

43 Mellone, S. et al. The Usefulness of a Targeted Next Generation Sequencing Gene Panel in Providing Molecular Diagnosis to Patients With a Broad Spectrum of Neurodevelopmental Disorders. Front Genet 13, 875182, doi:10.3389/fgene.2022.875182 (2022).

44 Christianson, A. L. et al. X linked severe mental retardation, craniofacial dysmorphology, epilepsy, ophthalmoplegia, and cerebellar atrophy in a large South African kindred is localised to Xq24-q27. J Med Genet 36, 759–766, doi:10.1136/jmg.36.10.759 (1999).

45 Ilie, A. et al. A potential gain-of-function variant of SLC9A6 leads to endosomal alkalinization and neuronal atrophy associated with Christianson Syndrome. Neurobiol Dis 121, 187–204, doi:10.1016/j.nbd.2018.10.002 (2019).

46 Okochi, R., Nihei, Y. & Ito, D. NDPACX: a newly defined X-linked Parkinsonian syndrome associated with SLC9A6 hemizygote mutation. Brain Commun 7, fcaf435, doi:10.1093/braincomms/fcaf435 (2025).

47 Yin, J. et al. Next Generation Sequencing of 134 Children with Autism Spectrum Disorder and Regression. Genes (Basel) 11, doi:10.3390/genes11080853 (2020).

48 Garbern, J. Y. et al. A mutation affecting the sodium/proton exchanger, SLC9A6, causes mental retardation with tau deposition. Brain 133, 1391–1402, doi:10.1093/brain/awq071 (2010).

49 Ilie, A., Weinstein, E., Boucher, A., McKinney, R. A. & Orlowski, J. Impaired posttranslational processing and trafficking of an endosomal Na+/H+ exchanger NHE6 mutant (Delta(370)WST(372)) associated with X-linked intellectual disability and autism. Neurochem Int 73, 192–203, doi:10.1016/j.neuint.2013.09.020 (2014).

50 Sarlus, H. & Heneka, M. T. Microglia in Alzheimer’s disease. J Clin Invest 127, 3240–3249, doi:10.1172/JCI90606 (2017).

51 Yanuck, S. F. Microglial Phagocytosis of Neurons: Diminishing Neuronal Loss in Traumatic, Infectious, Inflammatory, and Autoimmune CNS Disorders. Front Psychiatry 10, 712, doi:10.3389/fpsyt.2019.00712 (2019).

52 Mignot, C. et al. Novel mutation in SLC9A6 gene in a patient with Christianson syndrome and retinitis pigmentosum. Brain Dev 35, 172–176, doi:10.1016/j.braindev.2012.03.010 (2013).

53 Ishida, Y., Nayak, S., Mindell, J. A. & Grabe, M. A model of lysosomal pH regulation. J Gen Physiol 141, 705–720, doi:10.1085/jgp.201210930 (2013).

54 Xinhan, L. et al. Na+/H+ exchanger isoform 6 (NHE6/SLC9A6) is involved in clathrin-dependent endocytosis of transferrin. Am J Physiol Cell Physiol 301, C1431–1444, doi:10.1152/ajpcell.00154.2011 (2011).

55 Ohgaki, R. et al. The Na+/H+ exchanger NHE6 in the endosomal recycling system is involved in the development of apical bile canalicular surface domains in HepG2 cells. Mol Biol Cell 21, 1293–1304, doi:10.1091/mbc.e09-09-0767 (2010).

56 Lizarraga, S. B. et al. Human neurons from Christianson syndrome iPSCs reveal mutation-specific responses to rescue strategies. Sci Transl Med 13, doi:10.1126/scitranslmed.aaw0682 (2021).

57 Busa, W. B. & Nuccitelli, R. Metabolic regulation via intracellular pH. Am J Physiol 246, R409–438, doi:10.1152/ajpregu.1984.246.4.R409 (1984).

58 Munder, M. C. et al. A pH-driven transition of the cytoplasm from a fluid- to a solid-like state promotes entry into dormancy. Elife 5, doi:10.7554/eLife.09347 (2016).

59 Musgrove, E., Seaman, M. & Hedley, D. Relationship between cytoplasmic pH and proliferation during exponential growth and cellular quiescence. Exp Cell Res 172, 65–75, doi:10.1016/0014-4827(87)90093-0 (1987).

60 Casey, J. R., Grinstein, S. & Orlowski, J. Sensors and regulators of intracellular pH. Nat Rev Mol Cell Biol 11, 50–61, doi:10.1038/nrm2820 (2010).

61 Ouyang, Q. et al. Christianson syndrome protein NHE6 modulates TrkB endosomal signaling required for neuronal circuit development. Neuron 80, 97–112, doi:10.1016/j.neuron.2013.07.043 (2013).

62 Demuro, A., Parker, I. & Stutzmann, G. E. Calcium signaling and amyloid toxicity in Alzheimer disease. J Biol Chem 285, 12463–12468, doi:10.1074/jbc.R109.080895 (2010).

63 Ullman, J. C. et al. A mouse model of autism implicates endosome pH in the regulation of presynaptic calcium entry. Nat Commun 9, 330, doi:10.1038/s41467-017-02716-5 (2018).

64 Petitjean, H. et al. Loss of SLC9A6/NHE6 impairs nociception in a mouse model of Christianson syndrome. Pain 161, 2619–2628, doi:10.1097/j.pain.0000000000001961 (2020).

65 Deane, E. C. et al. Enhanced recruitment of endosomal Na+/H+ exchanger NHE6 into Dendritic spines of hippocampal pyramidal neurons during NMDA receptor-dependent long-term potentiation. J Neurosci 33, 595–610, doi:10.1523/JNEUROSCI.2583-12.2013 (2013).

66 Berridge, M. J., Bootman, M. D. & Roderick, H. L. Calcium signalling: dynamics, homeostasis and remodelling. Nat Rev Mol Cell Biol 4, 517–529, doi:10.1038/nrm1155 (2003).

67 Zhang, B. et al. Integrated systems approach identifies genetic nodes and networks in late-onset Alzheimer’s disease. Cell 153, 707–720, doi:10.1016/j.cell.2013.03.030 (2013).

68 Shimizu, H. et al. Crystal structure of an active form of BACE1, an enzyme responsible for amyloid beta protein production. Mol Cell Biol 28, 3663–3671, doi:10.1128/MCB.02185-07 (2008).

69 Olubiyi, O. O. & Strodel, B. Structures of the amyloid beta-peptides Abeta1-40 and Abeta1-42 as influenced by pH and a D-peptide. J Phys Chem B 116, 3280–3291, doi:10.1021/jp2076337 (2012).

70 Fernandez, M. A. et al. Loss of endosomal exchanger NHE6 leads to pathological changes in tau in human neurons. Stem Cell Reports 17, 2111–2126, doi:10.1016/j.stemcr.2022.08.001 (2022).

71 Shinohara, M., Tachibana, M., Kanekiyo, T. & Bu, G. Role of LRP1 in the pathogenesis of Alzheimer’s disease: evidence from clinical and preclinical studies. J Lipid Res 58, 1267–1281, doi:10.1194/jlr.R075796 (2017).

72 Truty, R. et al. Possible precision medicine implications from genetic testing using combined detection of sequence and intragenic copy number variants in a large cohort with childhood epilepsy. Epilepsia Open 4, 397–408, doi:10.1002/epi4.12348 (2019).

73 Fichou, Y. et al. Mutation in the SLC9A6 gene is not a frequent cause of sporadic Angelman-like syndrome. Eur J Hum Genet 17, 1378–1380, doi:10.1038/ejhg.2009.82 (2009).

74 Ouyang, Q. et al. Functional Assessment In Vivo of the Mouse Homolog of the Human Ala-9-Ser NHE6 Variant. eNeuro 6, doi:10.1523/ENEURO.0046-19.2019 (2019).

75 Li, Y., Xu, W., Mu, Y. & Zhang, J. Z. Acidic pH retards the fibrillization of human Islet Amyloid Polypeptide due to electrostatic repulsion of histidines. J Chem Phys 139, 055102, doi:10.1063/1.4817000 (2013).

76 Kondapalli, K. C. et al. A leak pathway for luminal protons in endosomes drives oncogenic signalling in glioblastoma. Nat Commun 6, 6289, doi:10.1038/ncomms7289 (2015).

77 Grabe, M., Wang, H. & Oster, G. The mechanochemistry of V-ATPase proton pumps. Biophys J 78, 2798–2813, doi:10.1016/S0006-3495(00)76823-8 (2000).

78 Marcoline, F. V., Ishida, Y., Mindell, J. A., Nayak, S. & Grabe, M. A mathematical model of osteoclast acidification during bone resorption. Bone 93, 167–180, doi:10.1016/j.bone.2016.09.007 (2016).

79 Lee, C. et al. A two-domain elevator mechanism for sodium/proton antiport. Nature 501, 573–577, doi:10.1038/nature12484 (2013).

80 Taglicht, D., Padan, E. & Schuldiner, S. Overproduction and purification of a functional Na+/H+ antiporter coded by nhaA (ant) from Escherichia coli. J Biol Chem 266, 11289–11294 (1991).

81 Winklemann, I. et al. Structure and elevator mechanism of the mammalian sodium/proton exchanger NHE9. EMBO J 39, e105908, doi:10.15252/embj.2020105908 (2020).

82 Rybak, S. L., Lanni, F. & Murphy, R. F. Theoretical considerations on the role of membrane potential in the regulation of endosomal pH. Biophys J 73, 674–687, doi:10.1016/S0006-3495(97)78102-5 (1997).

83 Ashkenazy, H. et al. ConSurf 2016: an improved methodology to estimate and visualize evolutionary conservation in macromolecules. Nucleic Acids Res 44, W344–350, doi:10.1093/nar/gkw408 (2016).

84 Kessel, A. & Ben-Tal, N. Free energy determinants of peptide association with lipid bilayers. Current topics in membranes 52, 205–253 (2002).

85 Gomez Ravetti, M., Rosso, O. A., Berretta, R. & Moscato, P. Uncovering molecular biomarkers that correlate cognitive decline with the changes of hippocampus’ gene expression profiles in Alzheimer’s disease. PLoS One 5, e10153, doi:10.1371/journal.pone.0010153 (2010).

