## Supplemental Information for "Endosomal pH Triggers Amyloid β Oligomerization and Maladaptive Phenotypic Plasticity in Alzheimer’s Disease"

[#]Equal contribution

Tables of Contents

• Supplementary Figures 1-7
• Supplementary Tables 1-2
• Supplementary References

**Supplementary Fig. 1: Cell type expression of endosomal $Na^{+}/H^{+}$ exchangers and pathway analysis of the NHE6 co-expression network.**

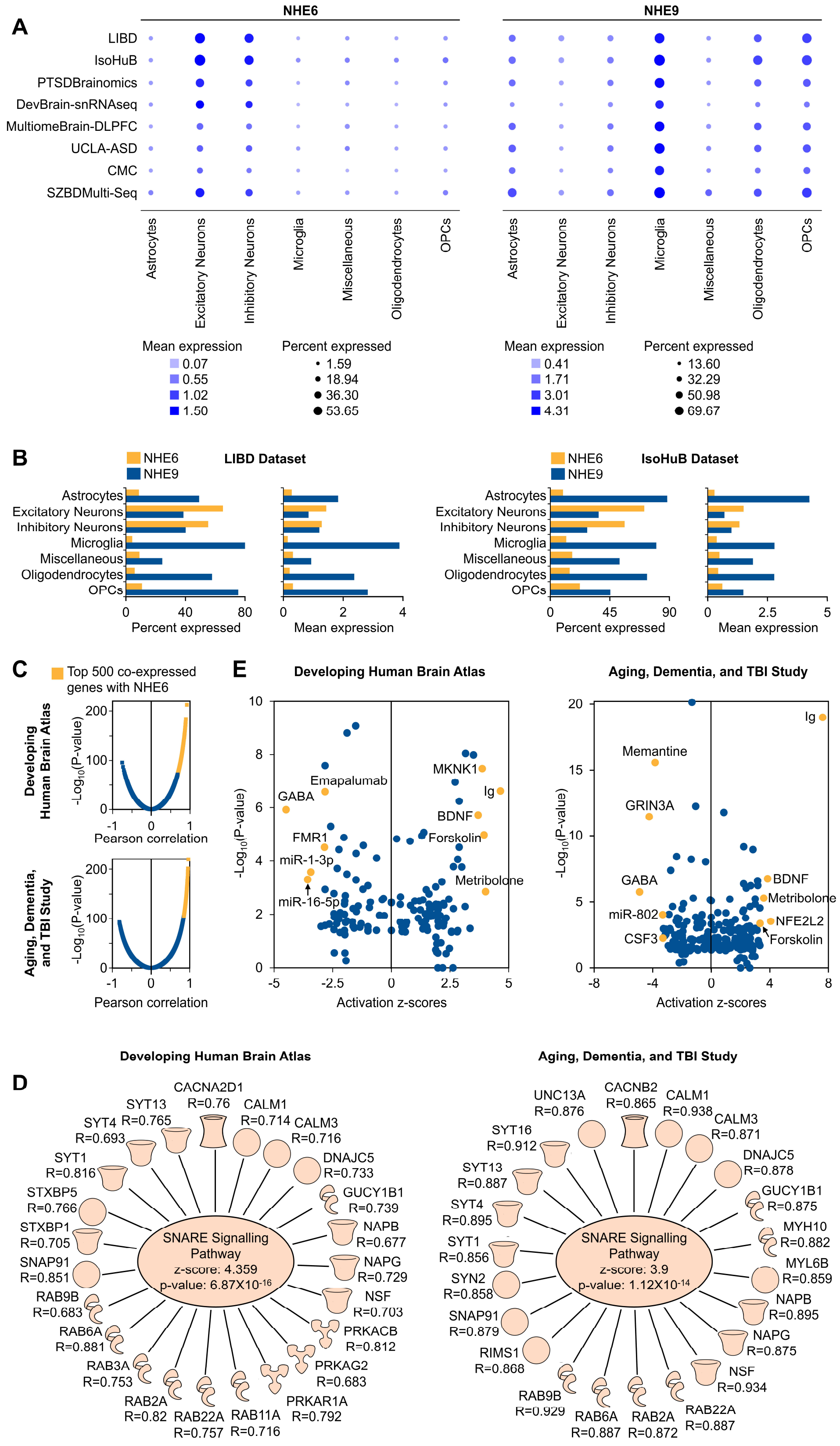

(A) Dot plot showing scaled mean expression (color) and percentage of cells expressing each gene (dot size) across annotated cell types from adult dorsolateral prefrontal cortex (DLPFC) from PsychENCODE Consortium data, including two normal datasets (LIBD, n=4; IsoHuB, n=10) and six disease datasets from patients with post-traumatic stress disorder (PTSD), major depressive disorder (MDD), autism spectrum disorder (ASD), Williams syndrome (WS), schizophrenia (SCZ), and bipolar disorder (BD), as well as matched controls: PTSDBrainomics (PTSD n=6, MDD n=4, control n=9), DevBrain-snRNAseq (ASD n=9, WS n=3, control n=4), MultiomeBrain-DLPFC (SCZ n=6, BD n=10, control n=5), UCLA-ASD (ASD n=27, control n=25), CMC (SCZ n=47, control n=53), and SZBDMulti-Seq (SCZ n=24, BD n=24, control n=24). (B) Bar plots showing NHE6 and NHE9 expression patterns in the LIBD (*left*) and IsoHuB (*right*) datasets, showing percent expressed and mean expression. Note that NHE6 expression is enriched in neuronal populations, including both excitatory and inhibitory subtypes, whereas NHE9 expression is enriched in glial populations, such as microglia and oligodendrocyte precursor cells (OPCs). (C) Scatter plots depicting correlation analysis of NHE6 expression with expression of (*top*) 52,375 transcripts across 524 brain samples from different regions (8 postconception weeks to 40 years of age) in the Developing Human Brain Atlas, and (*bottom*) 50,280 transcripts across 377 brain samples from different regions of aged donors with traumatic brain injury (TBI) and their matched controls from the Aging, Dementia, and TBI Study RNA-seq datasets. The x-axis indicates Pearson correlation, while the y-axis displays the negative log-base-10 of the p-value. The top 500 genes correlating with NHE6 are shown in yellow and were subjected to Ingenuity Pathway Analysis (IPA). (D) Network depicting the correlation (R) of NHE6 expression with genes in the SNARE signaling canonical pathway identified by IPA of the top 500 genes co-expressed with NHE6 in the normal Developing Human Brain Atlas (*left*) and the Aging, Dementia, and TBI Study (*right*) datasets, highlighting the role of NHE6 in vesicular trafficking and synaptic function. (E) Scatter plot depicting upstream regulators of NHE6 co-expressed genes identified by IPA of the top 500 genes co-expressed with NHE6 in the normal Developing Human Brain Atlas (*left*) and the Aging, Dementia, and TBI Study (*right*) datasets. The x-axis indicates z-score, while the y-axis displays the negative log-base-10 of the p-value. Top upstream regulators are colored yellow and labelled. Related to Figure 1.

**Supplementary Fig. 2: Structural and functional characterization of NHE6 patient variants.**

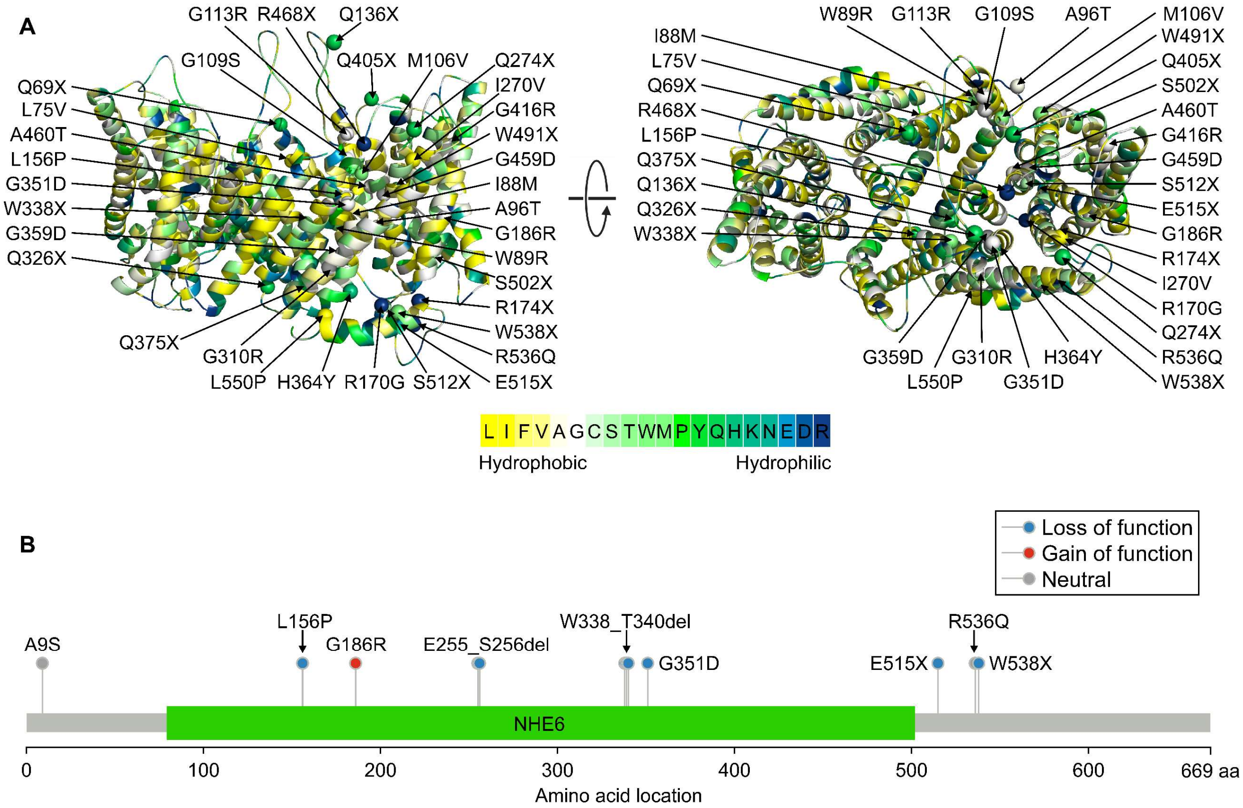


(A) Side (*left*) and top (*right*) views of the NHE6 dimer, including the transport domain and proximal C-terminal domain, based on the related NHE9 structure (PDB: 8PXB) and generated using the SWISS-MODEL server. The NHE6 model structure is colored according to hydrophobicity analysis using the blue-to-yellow color code, showing that the lipid-facing amino acids are overall hydrophobic, as expected. The hydrophobicity color bar is shown at the bottom. NHE6 missense and nonsense patient variants are depicted as α-carbon spheres on one monomer. (B) Lollipop representation summarizing functionally characterized NHE6 variants and their functional scoring. The membrane-embedded ion transport domain is indicated. Variants are categorized according to loss-of-function, neutral, or gain-of-function effects. Note that the two missense variants in the N-terminal segment (A9S) and C-terminal tail (R536Q) were neutral and may represent benign polymorphisms or have subtle functional effects, consistent with the fact that these regions do not participate directly in ion transport; however, truncation of the C-terminal tail (E515X and W538X) leads to loss of function, indicating that the C-terminal tail may have important regulatory roles. See also Supplementary Table 2. Related to Figure 2.

**Supplementary Fig. 3: Quantitative modelling of endosomal pH dynamics and amyloidogenic activity.**

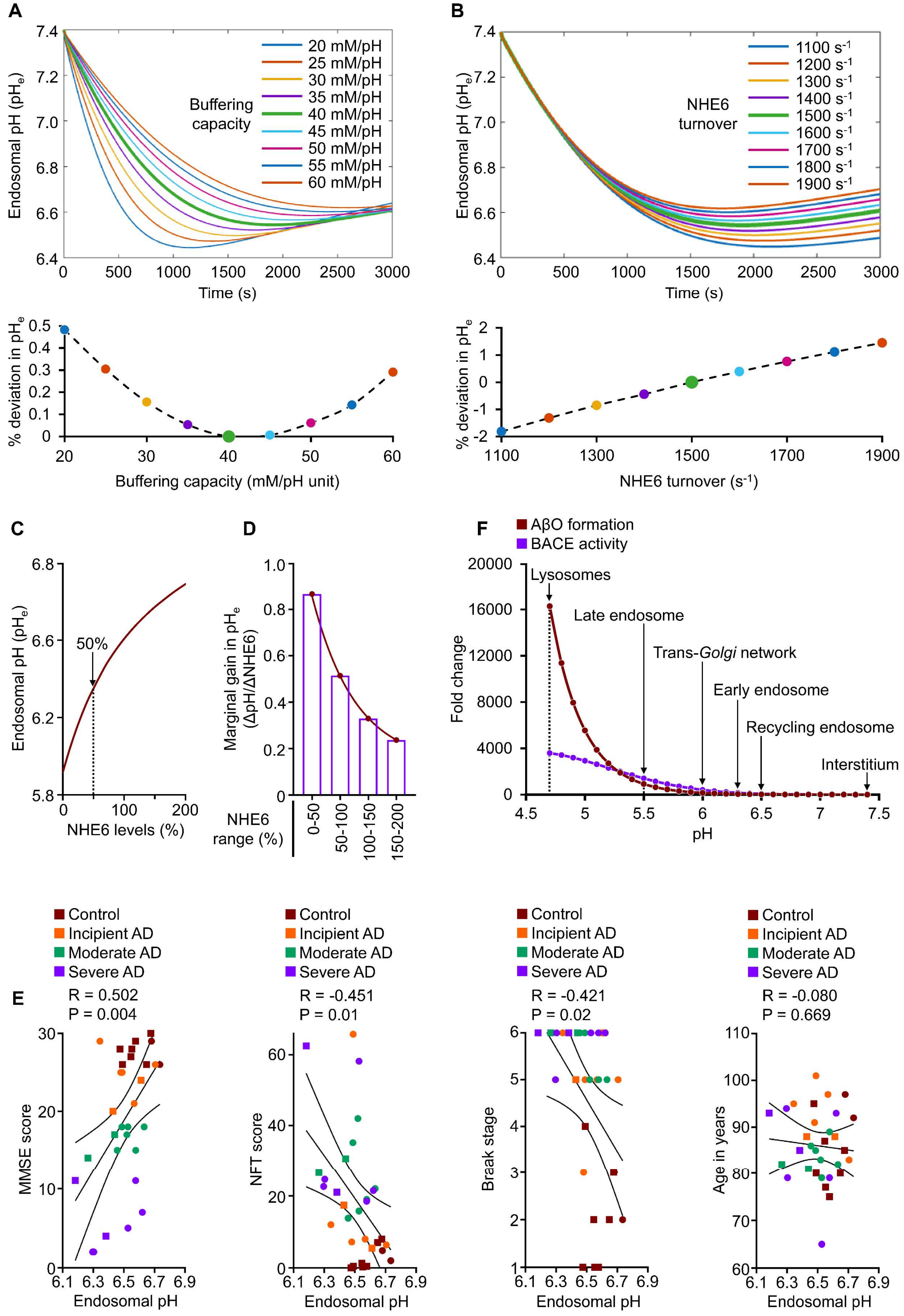

(A–B) Sensitivity analysis performed for buffering capacity (A, 20–60 mM/pH unit; baseline: 40 mM/pH unit) and NHE6 turnover rate (B, 1100–1900 ions/s; baseline: 1500 ions/s). Endosomal acidification dynamics are shown in the top panels, with the percentage deviation from baseline shown in the bottom panels. Buffering capacity primarily influenced the initial rate of acidification, whereas the NHE6 turnover rate predominantly affected steady-state pH. Deviations from baseline were minimal to modest (<0.5% and <2%, respectively), supporting the robustness of the model. (C) Plot showing NHE6 levels (percentage of baseline, ranging from 0–200%) versus steady-state endosomal pH (pHe) (y-axis), demonstrating a nonlinear saturable relationship. Note the apparent threshold-like behaviour below 50% of baseline expression, indicated by the dotted line, where the curve transitions from a steep pH dependence to a plateau. (D) Bar graph quantifying the diminishing marginal gain ($\Delta$pH/$\Delta$NHE6) in endosomal pH for sequential 50% increases in NHE6 expression, consistent with the threshold-like behaviour. (E) Scatter plots showing clinicopathological associations of estimated endosomal pH with antemortem cognitive function assessed by Mini-Mental State Examination (MMSE) score, and post-mortem neuropathology assessed by neurofibrillary tangle (NFT) score, Braak stage, and age in years, obtained from analysis of postmortem hippocampus (GSE1297) across disease stages (Control, Incipient AD, Moderate AD, Severe AD). Linear fit, Pearson correlation (R), and 95% confidence interval bands are shown. Each scatter point represents an individual postmortem brain; squares represent males and circles represent females. (F) Plot showing pH versus fold change of BACE activity and A$\beta$ oligomer formation (y-axis), illustrating nonlinear increase under increasingly acidic conditions across pH values ranging from interstitial pH (7.4) to lysosomal pH (4.7). Fold changes are relative to interstitial pH (7.4), with calculated values for different cellular compartments highlighted. Related to Figures 3 and 4.

**Supplementary Fig. 4: NHE6 co-expression with calcium signaling and synapse-related genes in the developing human brain.**

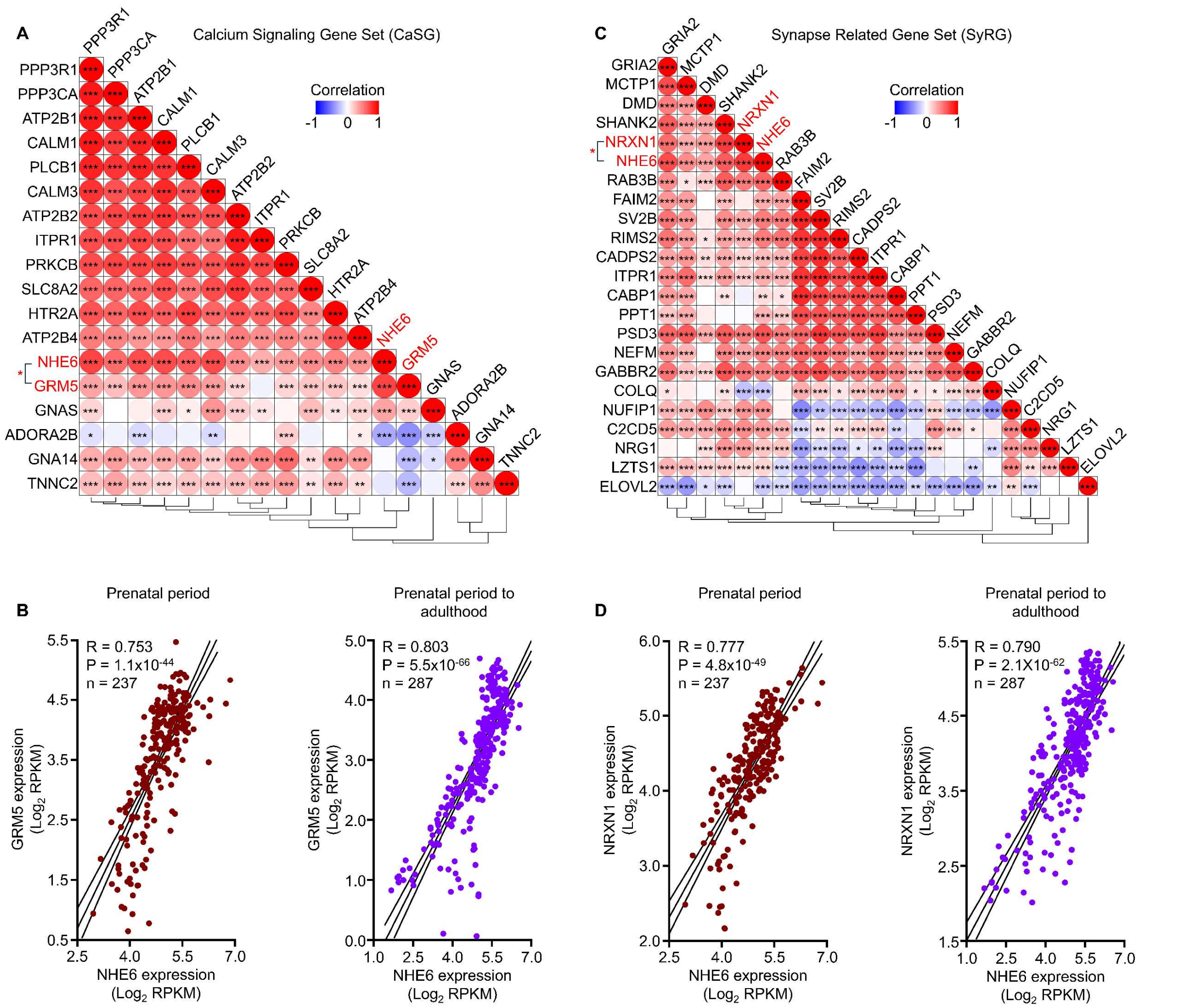


(A) Heatmap of the correlation coefficient matrix depicting associations between NHE6 and the 18-gene downregulated calcium signaling gene set in AD across the developing human brain. SLC25A6 was excluded because the dataset lacked values for this gene. Most genes show coordinated developmental expression, with NHE6 clustering alongside them. Strong co-expression was observed between NHE6 and the gene encoding metabotropic glutamate receptor 5 (GRM5) (*red asterisk*). (B) Scatter plots depicting the positive correlation of NHE6 with GRM5 in prenatal (*left*) and postnatal (*right*) development. (C) Heatmap of the correlation coefficient matrix depicting associations between NHE6 and the 22-gene downregulated synapse-related gene set in AD across the developing human brain. Most genes show coordinated developmental expression, with NHE6 clustering alongside them. Strong co-expression was observed between NHE6 and the gene encoding neurexin 1 (NRXN1) (*red asterisk*). (D) Scatter plots depicting the positive correlation of NHE6 with NRXN1 in prenatal (*left*) and postnatal (*right*) development. RNA sequencing gene expression data (8 postconception weeks to 40 years of age) were obtained from the BrainSpan atlas and are represented as log base 2 of reads per kilobase of exon model per million mapped reads (RPKM). $^{*}P < 0.05$, $^{**}P < 0.01$, $^{***}P < 0.001$ for each comparison in panels A and C. Linear fit, Pearson correlation (R), and 95% confidence interval bands are shown in B and D. Related to Figure 5.

**Supplementary Fig. 5: Relationship between NHE6 expression and the dysregulation of calcium signalling and synaptic dysfunction in Alzheimer's disease.**

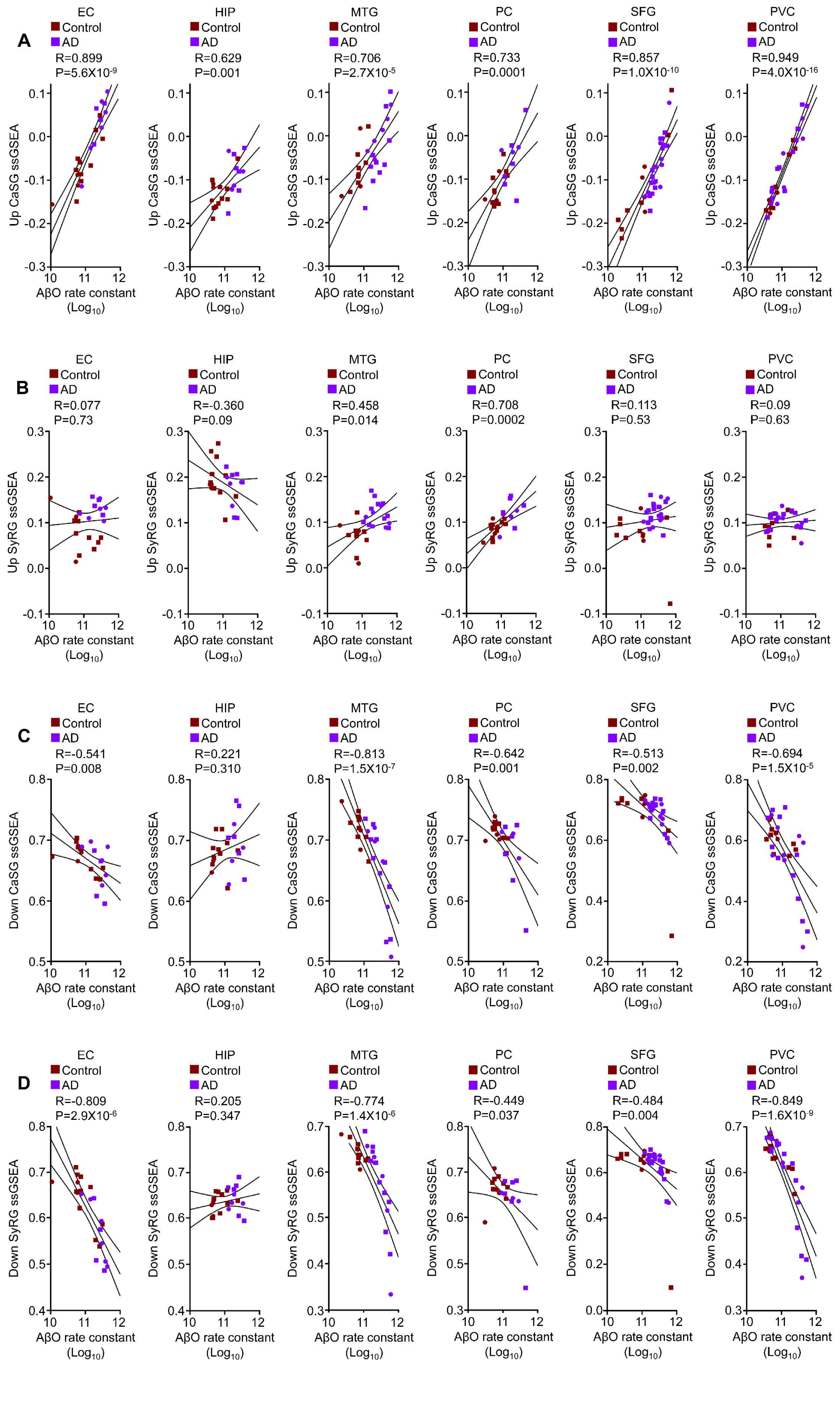

(A-B) Scatter plots depicting the correlation of the NHE6-associated AβO rate constant with ssGSEA scores for (A) the upregulated $Ca^{2+}$ signaling gene set (Up CaSG) and (B) the upregulated synapse-related gene set (Up SyRG) in control and AD across six brain regions: entorhinal cortex (EC), hippocampus (HIP), medial temporal gyrus (MTG), posterior cingulate (PC), superior frontal gyrus (SFG), and primary visual cortex (PVC). Note the positive association for ssGSEA scores of the upregulated $Ca^{2+}$ signaling gene set across all examined brain regions, whereas Up SyRG scores were either positively associated (in MTG and PC) or not significant (in all other regions). (C-D) Scatter plots depicting the correlation of the NHE6-associated AβO rate constant with ssGSEA scores for (C) the downregulated $Ca^{2+}$ signaling gene set (Down CaSG) and (D) the downregulated synapse-related gene set (Down SyRG) in control and AD across the same six brain regions. Note the negative association for Down CaSG and Down SyRG scores across brain regions, except for the hippocampus, which showed no significant association. Linear fit, Pearson correlation (R), and 95% confidence interval bands are shown. Data were obtained from analysis of postmortem brains from six brain regions (GSE5281). Each scatter point represents an individual postmortem brain; squares represent males and circles represent females. Related to Figure 5.

**Supplementary Fig. 6: Regional heterogeneity of NHE6-associated Aβ oligomerization in Alzheimer's disease.**

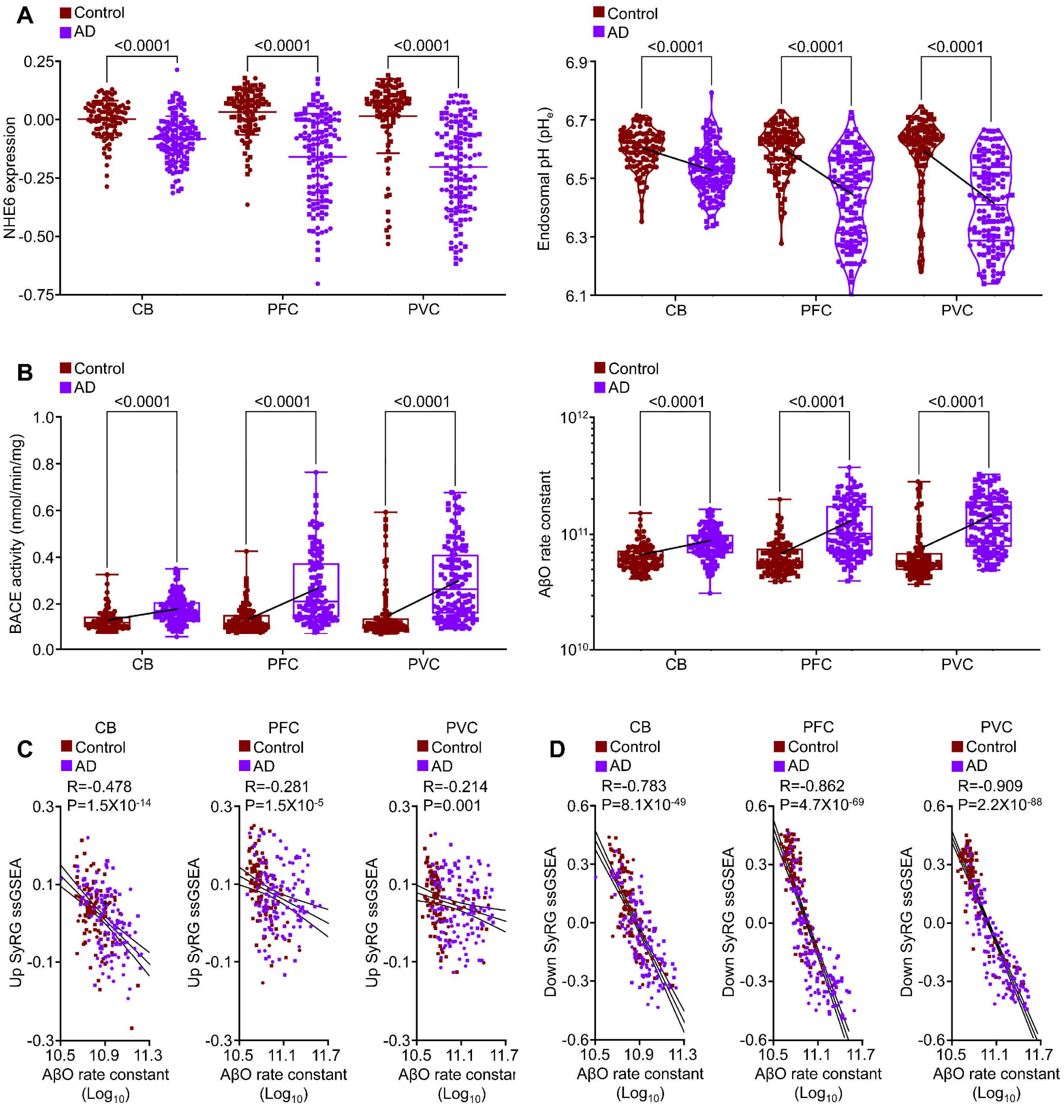


(A) Scatter plots with mean and standard deviation for normalized NHE6 expression (*left*) and violin plots with scatter showing estimated endosomal pH (*right*) in control and AD brains across three brain regions: cerebellum (CB), dorsolateral prefrontal cortex (PFC), and primary visual cortex (PVC). A line connecting the average pH values is overlaid. (B) Box-and-whisker plots showing specific BACE activity (*left*) and AβO rate constant (*right*), calculated from endosomal pH derived from NHE6 expression levels. A line connecting the average values is overlaid. P values in panels (A–B) were calculated by unpaired, two-tailed t test. Note the regional heterogeneity, with the least pronounced NHE6 downregulation, predicted endosomal hyperacidification, specific BACE activity, and AβO rate constant in the CB compared with the PFC and PVC. (C-D) Scatter plots showing correlations between the NHE6-associated AβO rate constant and ssGSEA scores for the (C) upregulated and (D) downregulated synapse-related gene sets (Up SyRG and Down SyRG) across the three brain regions: CB, PFC, and PVC. Linear fit, Pearson correlation (R), and 95% confidence interval bands are shown. Negative correlations were observed for both gene sets across all brain regions, with Down SyRG showing a stronger negative association with AβO rate constant than Up SyRG. Data are from the GSE44772 dataset. Each scatter point represents an individual postmortem brain; squares represent males and circles represent females. Related to Figure 5.

**Supplementary Fig. 7: NHE6-associated calcium signaling networks and structural determinants of pH-sensitive Aβ oligomerization.**

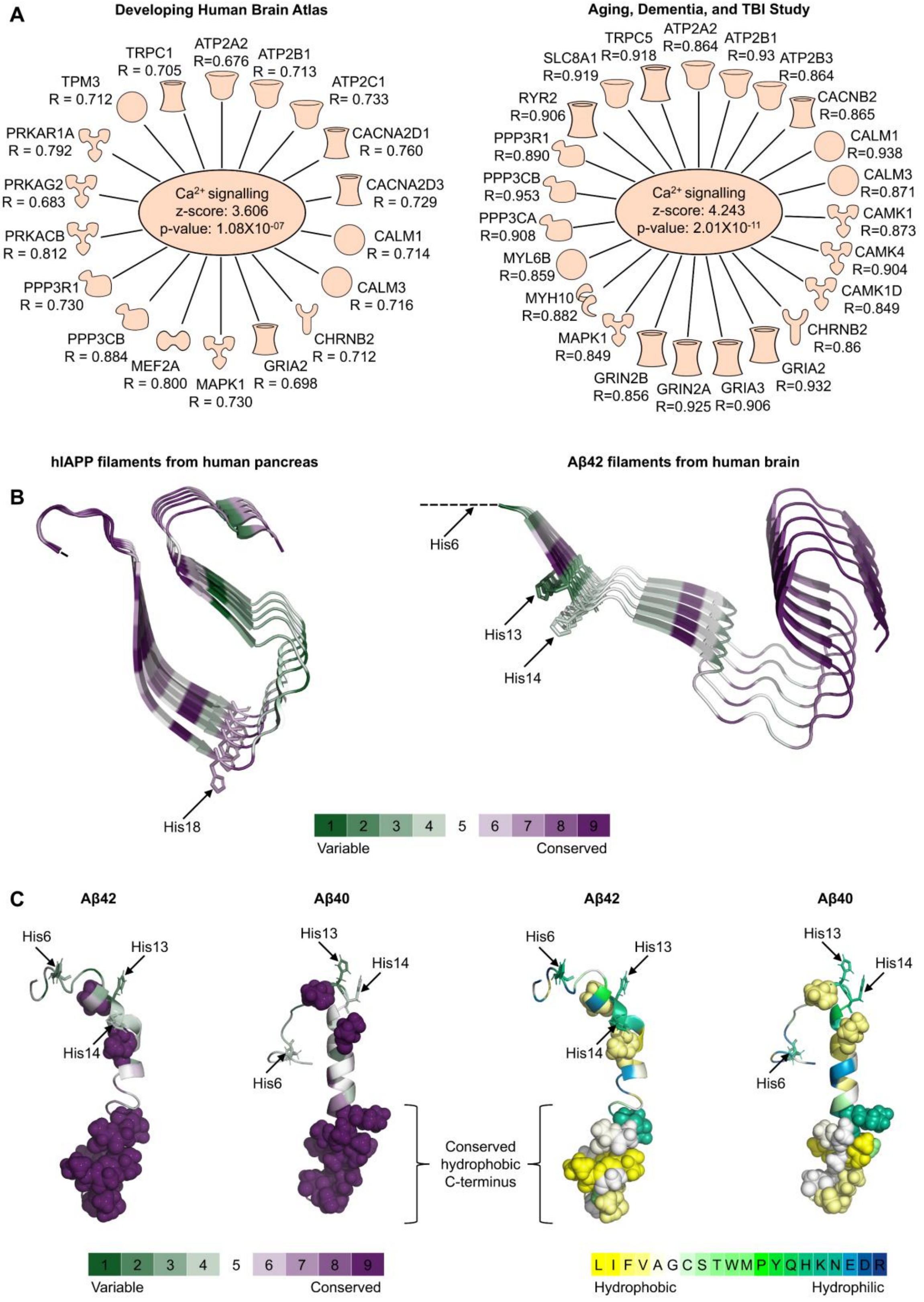

(A) Network depicting the correlation (R) of NHE6 expression with genes in the $Ca^{2+}$ signaling canonical pathway identified by Ingenuity Pathway Analysis (IPA) of the top 500 genes co-expressed with NHE6 in the normal Developing Human Brain Atlas (*left*) and the Aging, Dementia, and TBI Study (*right*) datasets, supporting a (patho)physiological relationship between endosomal pH regulation and calcium signaling. (B) Comparison of the structure of an hIAPP filament from human pancreas (PDB: 9ULZ) with the structure of an Aβ42 filament from the human brain (PDB: 7Q4B). Both are implicated in Alzheimer's disease and are known to cross-seed each other, yet show opposing effects to acidic pH. Structures are colored according to ConSurf evolutionary conservation scores using the green-to-purple color code, with histidines involved in pH effects shown as stick representations. (C) Aβ42 (PDB: 1Z0Q) and Aβ40 (PDB: 1BA4) monomer structures colored according to (*left*) ConSurf evolutionary conservation scores using the green-to-purple color code and (*right*) hydrophobicity analysis using the blue-to-yellow color code, with the color bar shown at the bottom. Highly conserved (ConSurf score 9) residues are displayed as space-filled atoms. Histidines (His 6, His 13, and His 14) implicated in pH-sensitive oligomerization are shown as stick representations; their protonation relieves electrostatic repulsion, allowing the conserved hydrophobic C-terminus (denoted by a brace) to drive oligomerization. Related to Discussion.

**Supplementary Table 1: Survey of genetic variants implicating NHE6 in a spectrum of neurological disorders**

| Variant type | Nucleotide change | Protein change | Location | CCS | dbSNP entry (ClinVar entry) | Functional prediction | Phenotypes | Ref. |
|---|---|---|---|---|---|---|---|---|
| Missense (27) | c.25G>T | p.A9S | NTS | 3 | rs201523857 (159933) | Polymorphism | Angelman-like syndrome | 1-3 |
| | c.157A>G | p.M53V | NTS | 5 | | Disease causing | ID | 4 |
| | c.168G>T | p.E56D | NTS | 6 | | Disease causing | Partial epilepsy without NDD | 5 |
| | c.171C>G | p.I57M | NTS | 1 | rs782296172 (379120) | Disease causing | ASD and regression | 6 |
| | c.223C>G | p.L75V | TM1 | 7 | rs782056346 (878535) | Disease causing | NDD | 7 |
| | c.264C>G | p.I88M | TM1 | 7 | | Disease causing | ASD | 8 |
| | c.265T>C | p.W89R | TM1 | 9 | | Disease causing | ID and atypical parkinsonism | 9 |
| | c.286G>A | p.A96T | TM1-2 loop | 4 | | Disease causing | ID | 10 |
| | c.316A>G | p.M106V | TM2 | 7 | | Disease causing | ID | 11 |
| | c.325G>A | p.G109S | TM2 | 9 | | Disease causing | CS | 12 |
| | c.337G>C | p.G113R | TM2 | 9 | | Disease causing | Partial epilepsy without NDD | 5 |
| | c.467T>C | p.L156P | TM3 | 9 | | Disease causing | ID | 2,13,14 |
| | c.508A>G | p.R170G | TM3-4 loop | 6 | rs796053280 (207238) | Disease causing | CS | 15 |
| | c.556G>A | p.G186R | TM4 | 9 | | Disease causing | CS | 16 |
| | c.808A>G | p.I270V | TM6 | 8 | | Disease causing | Hydrocephalus | 17 |
| | c.928G>A | p.G310R | TM7 | 8 | | Disease causing | CS | 18 |
| | c.1052G>A | p.G351D | TM8-9 loop | 9 | | Disease causing | CS and ESES | 2,14,19, 20 |
| | c.1076G>A | p.G359D | TM9 | 8 | rs796053285 | Disease causing | NDD and epilepsy | 21 |
| | c.1090C>T | p.H364Y | TM9 | 9 | | Disease causing | NDD | 22 |
| | c.1246G>A | p.G416R | TM11 | 4 | rs2521333666 (2502845) | Disease causing | ID | 23 |
| | c.1376G>A | p.G459D | TM12 | 8 | rs1569525357 (577815) | Disease causing | CS and ESES | 18,24 |
| | c.1378G>A | p.A460T | TM12 | 9 | | Disease causing | Epilepsy | 25 |
| | c.1607G>A | p.R536Q | CTD | 6 | rs146263125 (139207) | Polymorphism | ID, Schizophrenia | 26,27 |
| | c.1649T>C | p.L550P | CTD | 9 | rs796053287 | Disease causing | NDD and epilepsy, CS | 2,18,21 |
| | c.1735G>A | p.E579K | CTD | 4 | rs1556622379 (1006408) | Disease causing | ID, epilepsy, progressive brain atrophy, and large head | 28 |
| | c.1752G>T | p.L584F | CTD | 1 | | Disease causing | CS | 29 |
| | c.1777C>G | p.L593V | CTD | 4 | | Disease causing | ASD and regression | 6 |
| Nonsense (17) | c.190G>T | p.E64X | NTS | 4 | | Disease causing | CS and CBD | 19,30 |
| | c.205C>T | p.Q69X | NTS | 4 | | Disease causing | CS | 18 |
| | c.406C>T | p.Q136X | TM2-3 loop | 5 | | Disease causing | CS | 5 |
| | c.520C>T | p.R174X | TM4 | 6 | | Disease causing | Gross and fine motor delay, learning disability, autism, and seizures | 31 |
| | c.820C>T | p.Q274X | TM6-7 loop | 4 | | Disease causing | CS and retinitis pigmentosa | 32 |
| | c.976C>T | p.Q326X | TM8 | 7 | rs398124224 (95378) | Disease causing | ID | 33 |
| | c.1014G>A | p.W338X | TM8 | 9 | | Disease causing | CS and epilepsy | 34,35 |
| | c.1123C>T | p.Q375X | TM10 | 7 | | Disease causing | CS | 36 |

| | | | | | | | | |
|---|---|---|---|---|---|---|---|---|
| | c.1213C>T | p.Q405X | TM10-11 loop | 4 | | Disease causing | GDD/ID | 37 |
| | c.1402C>T | p.R468X | TM12-13 loop | 7 | rs122461162 (11477) | Disease causing | Angelman-like syndrome, CS, LGS, and epileptic encephalopathy | 5,19,36, 38-40 |
| | c.1472G>A | p.W491X | TM13 | 6 | | Disease causing | CS and LGS | 19,41 |
| | c.1473G>A | p.W491X | TM13 | 6 | | Disease causing | CS and ESES | 18,24 |
| | c.1505C>G | p.S502X | CTD | 4 | | Disease causing | NDD and early infantile epileptic encephalopathy | 42 |
| | c.1535C>G | p.S512X | CTD | 4 | | Disease causing | CS | 18 |
| | c.1543G>T | p.E515X | CTD | 4 | rs398123003 (92121) | Disease causing | ID, CS | 2,19,43 |
| | c.1614G>A | p.W538X | CTD | 7 | | Disease causing | CS and ESES | 2,19,44 |
| | c.1982T>G | p.L661X | CTD | 6 | rs587784399 (159932) | Disease causing | ID | 45 |
| Start-loss (2) | c.1A>G | p.M1? | NTS | 4 | rs782640388 (853911) | Disease causing | CS | 18 |
| | c.2T>G | p.M1? | NTS | 4 | rs1006154022 (383439) | Disease causing | Phenotype not reported | 46 |
| Splicing (20) | c.430-1G>A | | IVS2 as G-A -1 | | | | CS | 19,47 |
| | c.507+1G>A | | IVS3 ds G-A +1 | | | | CS | 18 |
| | c.507+1G>T | | IVS3 ds G-T +1 | | | | Generalized epilepsy | 48 |
| | c.508-1G>A | | IVS3 as G-A -1 | | rs797044508 (167702) | | CS and epilepsy | 18,49 |
| | c.508-1G>T | | IVS3 as G-T -1 | | | | CS | 18 |
| | c.584+1G>T | | IVS4 ds G-T +1 | | | | CS | 50 |
| | c.584+1G>A | | IVS4 ds G-A +1 | | rs796053282 | | CS | 18 |
| | c.584+3A>G | | IVS4 ds A-G +3 | | rs372679456 (805479) | | CS | 29 |
| | c.584+5G>A | | IVS4 ds G-A +5 | | rs796053284 (207242) | | CS, paediatric movement disorders, epilepsy | 18,21,51 ,52 |
| | c.697+1G>A | | IVS5 ds G-A +1 | | rs160319893 (625212) | | ID | 53,54 |
| | c.698-2A>G | | IVS5 as A-G -2 | | rs252121975 (2152353) | | Epileptic encephalopathy | 55 |
| | c.803+1G>A | | IVS6 ds G-A +1 | | rs155661745 (432837) | | CS, epilepsy, and GLUT1 deficiency syndrome | 18,24,56 ,57 |
| | c.946-1C>T | | IVS7 as C-T -1 | | rs149044510 (3371115) | | CS | 58 |
| | c.1052-2A>C | | IVS8 as A-C -2 | | | | Developmental disorder | 59 |
| | c.1141-8C>A | | IVS9 as C-A -8 | | | | CS | 60 |
| | c.1141-2A>G | | IVS9 as A-G -2 | | | | CS and ESES | 61 |
| | c.1255-1G>A | | IVS10 as G-A -1 | | | | CS and ESES | 62 |
| | c.1366+1G>C | | IVS11 ds G-C +1 | | rs2521368301 (2443320) | | CS | 63 |
| | c.1367-1G>A | | IVS11 as G-A -1 | | rs1603215383 (625179) | | CS | 64 |
| | c.1631+1G>A | | IVS14 ds G-A +1 | | rs796053283 (207241) | | CS, ID, and atlantoaxial instability | 65,66 |
| Regulatory (1) | c.*8A>T | | 3′ UTR | | rs200171451 (139210) | | Synesthesia | 67 |

| | | | | | | | | |
|---|---|---|---|---|---|---|---|---|
| Small insertions (10) | c.346dupC | p.L116Pfs*10 | TM2-3 loop | | rs796053293 (207251) | Disease causing | NDD and epilepsy | 21 |
| | c.444_451dupAGAAGTAT | p.F151* | TM3 | | | Disease causing | CS | 19 |
| | c.519dupT | p.R174Sfs*58 | TM4 | | | Disease causing | GDD and seizures | 68 |
| | c.534dupT | p.I179Yfs*53 | TM4 | | | Disease causing | CS | 18 |
| | c.676dupA | p.I226Nfs*6 | TM5 | | | Disease causing | CS | 69 |
| | c.1194dupG | p.L399Afs*12 | TM10 | | | Disease causing | CS and DEE | 70 |
| | c.1318dupA | p.R440Kfs*4 | TM11-12 loop | | | Disease causing | CS | 19 |
| | c.1409_1413dupCTGCC | p.T472Lfs*8 | TM12-13 loop | | | Disease causing | CS | 71 |
| | c.1464dupT | p.T489Yfs*23 | TM13 | | | Disease causing | CS and Parkinson's disease | 50,72 |
| | c.1550dupT | p.L517Ffs*5 | CTD | | | Disease causing | CS | 73 |
| Small deletions (30) | c.440delG# | p.S147Mfs*9 | TM2-3 loop | | rs2521144179 (3225530) | Disease causing | Angelman-like syndrome | 74 |
| | c.454_459delGAGTAT# | p.E152_Y153del | TM2-3 loop | | | Disease causing | ASD | 75 |
| | c.227_244delTCTTCATCCTGCTGCTCA | p.I76_L81del | TM1 | | | Disease causing | Epilepsy | 24 |
| | c.335_336delTG | p.V112Gfs*13 | TM2 | | | Disease causing | CS | 18 |
| | c.430-9_430-5delTTTTA | p.? | | | rs796053290 (207248) | | ID, CS | 76,77 |
| | c.477_481delCATAT | p.I160Lfs*5 | TM3 | | | Disease causing | CS and LGS | 40 |
| | c.488_501delCAGGTTATAGCCTG | p.A163Efs*64 | TM3 | | | Disease causing | Refractory epilepsy | 78 |
| | c.486_499delTGCAGGTTATAGCC | p.A163Efs*64 | TM3 | | | Disease causing | NDD/ID | 79 |
| | c.493delT | p.Y165Ifs*3 | TM3-4 loop | | | Disease causing | DEE | 80 |
| | c.507+1delGTAA | p.V144_R169del | TM2-3 loop to TM3-4 loop | | | | Angelman-like syndrome, ID | 26,38 |
| | c.512delA | p.H171Lfs*10 | TM3-4 loop | | | Disease causing | Developmental delay with seizures and movement disorder, CS | 72,81 |
| | c.512_513delAT | p.H171Lfs*60 | TM3-4 loop | | rs730882188 (11479) | Disease causing | Angelman-like syndrome, ID | 26,38 |
| | c.764_769delAAAGTG | p.E255_S256del | TM6 | | rs886037619 (11476) | Disease causing | Angelman-like syndrome | 38,82-84 |
| | c.803+1delG | p.? | | | | | CS | 18 |
| | c.803+3_803+6delAAGT | p.? | | | | | CS | 85 |
| | c.857_858delCA | p.T286Sfs*59 | TM6-7 loop | | rs796053297 (207255) | Disease causing | CS | 18 |
| | c.970_973delGAGT | p.E324Sfs*11 | TM7-8 loop | | | Disease causing | Developmental disorder | 86 |
| | c.976delC | p.Q326Sfs*10 | TM8 | | | Disease causing | ASD | 87 |
| | c.1013_1021delGGAGTACCT | p.W338_T340del | TM8 | | | Polymorphism | ID with tau deposition, CBD | 88-90 |

| | | | | | | | | |
|---|---|---|---|---|---|---|---|---|
| | c.1126_1130del CATAG | p.H376Nfs *2 | TM10 | | | Disease causing | CS, early infantile epileptic encephalopathy | 18,81 |
| | c.1181_1183del TCT | p.F394del | TM10 | | | Disease causing | DEE and ESES | 91 |
| | c.1203_1204del GT | p.F402Hfs* 8 | TM10 | | rs1064793575 (418996) | Disease causing | NDD and epilepsy | 21 |
| | c.1306_1309del CTTA | p.L436Ifs* 15 | TM11-12 loop | | rs2071147098 (949999) | Disease causing | CS | 5,18 |
| | c.1357_1359del ATG | p.M453del | TM12 | | | Disease causing | PD | 92 |
| | c.1450delC | p.L484* | TM13 | | | Disease causing | GDD/ID | 37 |
| | c.1458_1459del GT | p.F488Yfs* 23 | TM13 | | | Disease causing | ID | 53 |
| | c.1481delG | p.G494Vfs *53 | TM13 | | | Disease causing | ID | 93 |
| | c.1569_1573del AAGGA | p.R524Nfs *17 | CTD | | rs1603219805 (807688) | Disease causing | Dystonia | 94,95 |
| | c.1595_1613del CTTGGCTTT TCCGGATGT G | p.A532Gfs *9 | CTD | | (3362567) | Disease causing | CS | 29 |
| | c.1632-19_1632-3del17 | p.? | | | | | CS | 96 |
| Small indels (3) | c.742_743delC TinsG | p.L248Afs* 17 | TM6 | | | Disease causing | Epileptic encephalopathy | 55 |
| | c.1013_1025del 13insTCAGCC | p.W338Ffs *13 | TM8 | | | Disease causing | Epilepsy | 24 |
| | c.1284_1293del 10ins GTCTTGGGA AGACT | p.N428Kfs *11 | TM11 | | | Disease causing | CS | 18 |
| Gross deletions (10) | 120.7 kb including intron 10 to exon 16 and 3′-UTR (c.1237–556_*92 045del) | | | | | | CS | 18,19 |
| | 40 Mb including exon 5-14 | | | | | | CS and ESES | 97 |
| | 2073 bp including exon 1 and some of intron 1; (c.−1296_325+ 453 del) | | | | | | CS | 18 |
| | 314 kb including exon 15-16 of NHE6 and FHL1, MAP7D3 & GPR112 genes | | | | | | CS | 98 |
| | 336 bp including exon 1 | | | | | | ID | 99 |
| | 38 bp, c.316_325+28del | | | | | | Intractable early-onset epilepsy, DEE | 80,100 |
| | 4722 bp | | | | | | CS | 101 |
| | 9 kb partial | | | | | | Severe ID | 102 |
| | including entire gene | | | | | | Epilepsy | 24 |
| | including exon 4-7 | | | | | | Hereditary ataxia | 103 |

This overview compiles studies linking NHE6 genetic variants to various neurological disorders. Amino acids are denoted using their single-letter codes in protein sequences. Evolutionary conservation of mutated residues was assessed with ConSurf analysis, assigning scores from 1 (most variable) to 9 (invariant), whereas functional predictions were generated using the MutationTaster algorithm. Transcript positions are based on reference sequence NM_006359; variants that occur specifically in the longer NHE6 isoform (marked with #) are instead based on NM_001042537. Residue locations for structural features are assigned based on alignment with the structure of the related endosomal NHE9 (PDB: 8PXB). Abbreviations: 3′ UTR, 3-prime untranslated region; as, acceptor splice site; ASD, autism spectrum disorder; CBD, corticobasal degeneration; CCS, ConSurf conservation score; CS, Christianson syndrome; CTD, C-terminal domain; DEE, developmental and epileptic encephalopathy; del, deletion; ds, donor splice site; dup, duplication; ESES, electrical status epilepticus during slow-wave sleep; fs, frameshift; GDD, global developmental delay; GLUT1, glucose transporter type 1; ID, intellectual disability; ins, insertion; IVS, intervening sequence (intron); LGS, Lennox-Gastaut syndrome; NDD, neurodevelopmental disorder; NTS, N-terminal segment; PD, Parkinson's disease; Ref., reference; TM, transmembrane segment; X, stop codon. Related to Figure 2.

**Supplementary Table 2: Model parameters**

| Parameter description | Value | Units | Ref. |
|---|---|---|---|
| Endosome radius | 0.35 | µm | 104,105 |
| Endosome volume | $1.80X10^{-16}$ | L | |
| Endosome surface area | $1.54X10^{-8}$ | $cm^2$ | |
| Endosomal pH in wild type neurons (steady state) | 6.57 | | 106 |
| Endosomal pH in NHE6 null neurons (steady state) | 5.88 | | |
| NHE turnover | 1500 | ions/s | 107 |
| Luminal pH (initial) | 7.4 | | 108 |
| Luminal sodium concentration (initial) | 145 | mM | |
| Luminal potassium concentration (initial) | 5 | mM | |
| Luminal chloride concentration (initial) | 110 | mM | |
| Cytosolic pH | 7.2 | | |
| Cytosolic sodium concentration | 10 | mM | |
| Cytosolic potassium concentration | 145 | mM | |
| Cytosolic chloride concentration | 10 | mM | |
| Sodium permeability | $9.6x10^{-7}$ | cm/s | 109 |
| Potassium permeability | $7.1x10^{-7}$ | cm/s | |
| Chloride permeability | $1.2x10^{-5}$ | cm/s | |
| Surface potential | mV | -50 | 108 |
| Bilayer capacitance | 1 | $\mu F/cm^2$ | |
| Proton permeability | $6x10^{-5}$ | cm/s | |
| Buffering capacity | 40 | mM/pH | |
| Osmotic coefficient | 0.73 | | |
| Partial molar volume of water | 18 | $cm^3$ /mol | 110 |
| Cytoplasmic osmolyte concentration | 290 | mM | |

**Supplementary References:**


1 Fichou, Y. *et al.* Mutation in the SLC9A6 gene is not a frequent cause of sporadic Angelman-like syndrome. *Eur J Hum Genet* **17**, 1378-1380, doi:10.1038/ejhg.2009.82 (2009).

2 Ilie, A. *et al.* Assorted dysfunctions of endosomal alkali cation/proton exchanger SLC9A6 variants linked to Christianson syndrome. *J Biol Chem* **295**, 7075-7095, doi:10.1074/jbc.RA120.012614 (2020).

3 Ouyang, Q. *et al.* Functional Assessment In Vivo of the Mouse Homolog of the Human Ala-9-Ser NHE6 Variant. *eNeuro* **6**, doi:10.1523/ENEURO.0046-19.2019 (2019).

4 Ghalamkari, S., Mianesaz, H., Chitsaz, A., Ghazavi, M. & Salehi, M. Proband-Only Exome Sequencing for Intellectual Disability in Iran: Diagnostic Yield and Genetic Insights. *Am J Med Genet A* **197**, e63915, doi:10.1002/ajmg.a.63915 (2025).

5 Jiao, J. P. *et al.* Missense variants in SLC9A6 cause partial epilepsy without neurodevelopmental delay. *Orphanet J Rare Dis* **20**, 380, doi:10.1186/s13023-025-03924-9 (2025).

6 Yin, J. *et al.* Next Generation Sequencing of 134 Children with Autism Spectrum Disorder and Regression. *Genes (Basel)* **11**, doi:10.3390/genes11080853 (2020).

7 Pranav Chand, R. *et al.* Proband only exome sequencing in 403 Indian children with neurodevelopmental disorders: Diagnostic yield, utility and challenges in a resource-limited setting. *Eur J Med Genet* **66**, 104730, doi:10.1016/j.ejmg.2023.104730 (2023).

8 Mellone, S. *et al.* The Usefulness of a Targeted Next Generation Sequencing Gene Panel in Providing Molecular Diagnosis to Patients With a Broad Spectrum of Neurodevelopmental Disorders. *Front Genet* **13**, 875182, doi:10.3389/fgene.2022.875182 (2022).

9 Nan, H. *et al.* Novel SLC9A6 Variation in Female Carriers With Intellectual Disability and Atypical Parkinsonism. *Neurol Genet* **8**, e651, doi:10.1212/NXG.0000000000000651 (2022).

10 Wang, J., Wang, Y., Wang, L., Chen, W. Y. & Sheng, M. The diagnostic yield of intellectual disability: combined whole genome low-coverage sequencing and medical exome sequencing. *BMC Med Genomics* **13**, 70, doi:10.1186/s12920-020-0726-x (2020).

11 Ibarluzea, N. *et al.* Targeted Next-Generation Sequencing in Patients with Suggestive X-Linked Intellectual Disability. *Genes (Basel)* **11**, doi:10.3390/genes11010051 (2020).

12 Heron, D. *et al.* A large cohort study of prenatal exome sequencing redefines diagnosis in fetal corpus callosum anomalies. *Brain* **148**, 4253-4258, doi:10.1093/brain/awaf311 (2025).

13 Hu, H. *et al.* Mutation screening in 86 known X-linked mental retardation genes by droplet-based multiplex PCR and massive parallel sequencing. *Hugo J* **3**, 41-49, doi:10.1007/s11568-010-9137-y (2009).

14 Prasad, H. & Rao, R. Amyloid clearance defect in ApoE4 astrocytes is reversed by epigenetic correction of endosomal pH. *Proc Natl Acad Sci U S A* **115**, E6640-E6649, doi:10.1073/pnas.1801612115 (2018).

15 Ouyang, X. *et al.* Clinical Utility of Rapid Exome Sequencing Combined With Mitochondrial DNA Sequencing in Critically Ill Pediatric Patients With Suspected Genetic Disorders. *Front Genet* **12**, 725259, doi:10.3389/fgene.2021.725259 (2021).

16 Ilie, A. *et al.* A potential gain-of-function variant of SLC9A6 leads to endosomal alkalinization and neuronal atrophy associated with Christianson Syndrome. *Neurobiol Dis* **121**, 187-204, doi:10.1016/j.nbd.2018.10.002 (2019).

17 Weitensteiner, V. *et al.* Exome sequencing in syndromic brain malformations identifies novel mutations in ACTB, and SLC9A6, and suggests BAZ1A as a new candidate gene. *Birth Defects Res* **110**, 587-597, doi:10.1002/bdr2.1200 (2018).

18 Kavanaugh, B. C. *et al.* Christianson syndrome across the lifespan: genetic mutations and longitudinal study in children, adolescents, and adults. *J Med Genet* **61**, 1031-1039, doi:10.1136/jmg-2024-109973 (2024).

19 Pescosolido, M. F. *et al.* Genetic and phenotypic diversity of NHE6 mutations in Christianson syndrome. *Ann Neurol* **76**, 581-593, doi:10.1002/ana.24225 (2014).

20 Lizarraga, S. B. *et al.* Human neurons from Christianson syndrome iPSCs reveal mutation-specific responses to rescue strategies. *Sci Transl Med* **13**, doi:10.1126/scitranslmed.aaw0682 (2021).
21 Lindy, A. S. *et al.* Diagnostic outcomes for genetic testing of 70 genes in 8565 patients with epilepsy and neurodevelopmental disorders. *Epilepsia* **59**, 1062-1071, doi:10.1111/epi.14074 (2018).
22 Wang, T. *et al.* Targeted sequencing and integrative analysis of 3,195 Chinese patients with neurodevelopmental disorders prioritized 26 novel candidate genes. *J Genet Genomics* **48**, 312-323, doi:10.1016/j.jgg.2021.03.002 (2021).
23 Hussain, S. I. *et al.* Structural and functional implications of SLC13A3 and SLC9A6 mutations: an in silico approach to understanding intellectual disability. *BMC Neurol* **23**, 353, doi:10.1186/s12883-023-03397-y (2023).
24 Truty, R. *et al.* Possible precision medicine implications from genetic testing using combined detection of sequence and intragenic copy number variants in a large cohort with childhood epilepsy. *Epilepsia Open* **4**, 397-408, doi:10.1002/epi4.12348 (2019).
25 Gall, K. *et al.* Next-generation sequencing in childhood-onset epilepsies: Diagnostic yield and impact on neuronal ceroid lipofuscinosis type 2 (CLN2) disease diagnosis. *PLoS One* **16**, e0255933, doi:10.1371/journal.pone.0255933 (2021).
26 Tarpey, P. S. *et al.* A systematic, large-scale resequencing screen of X-chromosome coding exons in mental retardation. *Nat Genet* **41**, 535-543, doi:10.1038/ng.367 (2009).
27 Piton, A. *et al.* Systematic resequencing of X-chromosome synaptic genes in autism spectrum disorder and schizophrenia. *Mol Psychiatry* **16**, 867-880, doi:10.1038/mp.2010.54 (2011).
28 Padmanabha, H., Saini, A. G., Sahu, J. K. & Singhi, P. Syndrome of X linked intellectual disability, epilepsy, progressive brain atrophy and large head associated with SLC9A6 mutation. *BMJ Case Rep* **2017**, doi:10.1136/bcr-2017-222050 (2017).
29 Mir, A. *et al.* SLC gene mutations and pediatric neurological disorders: diverse clinical phenotypes in a Saudi Arabian population. *Hum Genet* **141**, 81-99, doi:10.1007/s00439-021-02404-x (2022).
30 Sinajon, P., Verbaan, D. & So, J. The expanding phenotypic spectrum of female SLC9A6 mutation carriers: a case series and review of the literature. *Hum Genet* **135**, 841-850, doi:10.1007/s00439-016-1675-5 (2016).
31 Monies, D. *et al.* Lessons Learned from Large-Scale, First-Tier Clinical Exome Sequencing in a Highly Consanguineous Population. *Am J Hum Genet* **104**, 1182-1201, doi:10.1016/j.ajhg.2019.04.011 (2019).
32 Mignot, C. *et al.* Novel mutation in SLC9A6 gene in a patient with Christianson syndrome and retinitis pigmentosum. *Brain Dev* **35**, 172-176, doi:10.1016/j.braindev.2012.03.010 (2013).
33 LaDuca, H. *et al.* Exome sequencing covers >98% of mutations identified on targeted next generation sequencing panels. *PLoS One* **12**, e0170843, doi:10.1371/journal.pone.0170843 (2017).
34 Zhao, X. *et al.* Genetic analysis and identification of novel variations in Chinese patients with pediatric epilepsy by whole-exome sequencing. *Neurol Sci* **43**, 4439-4451, doi:10.1007/s10072-022-05953-9 (2022).
35 Peng, X. *et al.* [Clinical features and genetic analysis of a child with Christianson syndrome due to variant of SLC9A6 gene]. *Zhonghua Yi Xue Yi Chuan Xue Za Zhi* **42**, 411-418, doi:10.3760/cma.j.cn511374-20240919-00499 (2025).
36 Schroer, R. J. *et al.* Natural history of Christianson syndrome. *Am J Med Genet A* **152A**, 2775-2783, doi:10.1002/ajmg.a.33093 (2010).
37 Lin, L. *et al.* Clinical and genetic characteristics and prenatal diagnosis of patients presented GDD/ID with rare monogenic causes. *Orphanet J Rare Dis* **15**, 317, doi:10.1186/s13023-020-01599-y (2020).
38 Gilfillan, G. D. *et al.* SLC9A6 mutations cause X-linked mental retardation, microcephaly, epilepsy, and ataxia, a phenotype mimicking Angelman syndrome. *Am J Hum Genet* **82**, 1003-1010, doi:10.1016/j.ajhg.2008.01.013 (2008).
39 Hamdan, F. F. *et al.* High Rate of Recurrent De Novo Mutations in Developmental and Epileptic Encephalopathies. *Am J Hum Genet* **101**, 664-685, doi:10.1016/j.ajhg.2017.09.008 (2017).

40 Ikeda, A. *et al.* Epilepsy in Christianson syndrome: Two cases of Lennox-Gastaut syndrome and a review of literature. *Epilepsy Behav Rep* **13**, 100349, doi:10.1016/j.ebr.2019.100349 (2020).

41 Ream, M. A. & Mikati, M. A. Clinical utility of genetic testing in pediatric drug-resistant epilepsy: a pilot study. *Epilepsy Behav* **37**, 241-248, doi:10.1016/j.yebeh.2014.06.018 (2014).

42 Fernandez-Marmiesse, A. *et al.* Rare Variants in 48 Genes Account for 42% of Cases of Epilepsy With or Without Neurodevelopmental Delay in 246 Pediatric Patients. *Front Neurosci* **13**, 1135, doi:10.3389/fnins.2019.01135 (2019).

43 Schuurs-Hoeijmakers, J. H. *et al.* Identification of pathogenic gene variants in small families with intellectually disabled siblings by exome sequencing. *J Med Genet* **50**, 802-811, doi:10.1136/jmedgenet-2013-101644 (2013).

44 Coorg, R. & Weisenberg, J. L. Successful Treatment of Electrographic Status Epilepticus of Sleep With Felbamate in a Patient With SLC9A6 Mutation. *Pediatr Neurol* **53**, 527-531, doi:10.1016/j.pediatrneurol.2015.07.007 (2015).

45 Tan, C. A. *et al.* Characterization of patients referred for non-specific intellectual disability testing: the importance of autosomal genes for diagnosis. *Clin Genet* **89**, 478-483, doi:10.1111/cge.12575 (2016).

46 Capalbo, A. *et al.* Optimizing clinical exome design and parallel gene-testing for recessive genetic conditions in preconception carrier screening: Translational research genomic data from 14,125 exomes. *PLoS Genet* **15**, e1008409, doi:10.1371/journal.pgen.1008409 (2019).

47 Bosemani, T. *et al.* Christianson syndrome: spectrum of neuroimaging findings. *Neuropediatrics* **45**, 247-251, doi:10.1055/s-0033-1363091 (2014).

48 Leduc-Pessah, H., White-Brown, A., Hartley, T., Pohl, D. & Dyment, D. A. The Benefit of Multigene Panel Testing for the Diagnosis and Management of the Genetic Epilepsies. *Genes (Basel)* **13**, doi:10.3390/genes13050872 (2022).

49 Butler, K. M., da Silva, C., Alexander, J. J., Hegde, M. & Escayg, A. Diagnostic Yield From 339 Epilepsy Patients Screened on a Clinical Gene Panel. *Pediatr Neurol* **77**, 61-66, doi:10.1016/j.pediatrneurol.2017.09.003 (2017).

50 Riess, A. *et al.* Novel SLC9A6 mutations in two families with Christianson syndrome. *Clin Genet* **83**, 596-597, doi:10.1111/j.1399-0004.2012.01948.x (2013).

51 Cordeiro, D. *et al.* Genetic landscape of pediatric movement disorders and management implications. *Neurol Genet* **4**, e265, doi:10.1212/NXG.0000000000000265 (2018).

52 Mercimek-Mahmutoglu, S. *et al.* Diagnostic yield of genetic testing in epileptic encephalopathy in childhood. *Epilepsia* **56**, 707-716, doi:10.1111/epi.12954 (2015).

53 Grozeva, D. *et al.* Targeted Next-Generation Sequencing Analysis of 1,000 Individuals with Intellectual Disability. *Hum Mutat* **36**, 1197-1204, doi:10.1002/humu.22901 (2015).

54 Sanchis-Juan, A. *et al.* Rare Genetic Variation in 135 Families With Family History Suggestive of X-Linked Intellectual Disability. *Front Genet* **10**, 578, doi:10.3389/fgene.2019.00578 (2019).

55 Fung, C. W., Kwong, A. K. & Wong, V. C. Gene panel analysis for nonsyndromic cryptogenic neonatal/infantile epileptic encephalopathy. *Epilepsia Open* **2**, 236-243, doi:10.1002/epi4.12055 (2017).

56 Petraityte, G. *et al.* Donor Splice Site Variant in SLC9A6 Causes Christianson Syndrome in a Lithuanian Family: A Case Report. *Medicina (Kaunas)* **58**, doi:10.3390/medicina58030351 (2022).

57 Sanchez-Lijarcio, O. *et al.* The clinical and biochemical hallmarks generally associated with GLUT1DS may be caused by defects in genes other than SLC2A1. *Clin Genet* **102**, 40-55, doi:10.1111/cge.14138 (2022).

58 Yang, L. *et al.* Use of medical exome sequencing for identification of underlying genetic defects in NICU: Experience in a cohort of 2303 neonates in China. *Clin Genet* **101**, 101-109, doi:10.1111/cge.14075 (2022).

59 Deciphering Developmental Disorders Study. Large-scale discovery of novel genetic causes of developmental disorders. *Nature* **519**, 223-228, doi:10.1038/nature14135 (2015).

60 Ieda, D. *et al.* A novel splicing mutation in SLC9A6 in a boy with Christianson syndrome. *Hum Genome Var* **6**, 15, doi:10.1038/s41439-019-0046-x (2019).
61 Liu, X., Xie, L., Fang, Z. & Jiang, L. Case Report: Novel SLC9A6 Splicing Variant in a Chinese Boy With Christianson Syndrome With Electrical Status Epilepticus During Sleep. *Front Neurol* **12**, 796283, doi:10.3389/fneur.2021.796283 (2021).
62 Zanni, G. *et al.* A novel mutation in the endosomal Na+/H+ exchanger NHE6 (SLC9A6) causes Christianson syndrome with electrical status epilepticus during slow-wave sleep (ESES). *Epilepsy Res* **108**, 811-815, doi:10.1016/j.eplepsyres.2014.02.009 (2014).
63 Dong, Y. *et al.* Clinical and genetic analysis of Christianson syndrome caused by variant of SLC9A6: case report and literature review. *Front Neurol* **14**, 1152696, doi:10.3389/fneur.2023.1152696 (2023).
64 Zhang, X. *et al.* Christianson syndrome: A novel splicing variant of SLC9A6 causes exon skipping in a Chinese boy and a literature review. *J Clin Lab Anal* **36**, e24123, doi:10.1002/jcla.24123 (2022).
65 McSherry, M. *et al.* Identification of candidate gene FAM183A and novel pathogenic variants in known genes: High genetic heterogeneity for autosomal recessive intellectual disability. *PLoS One* **13**, e0208324, doi:10.1371/journal.pone.0208324 (2018).
66 Guven, N. E. *et al.* Atlantoaxial Instability due to Os Odontoideum in a Child with Christianson Syndrome. *Mol Syndromol* **15**, 398-402, doi:10.1159/000538015 (2024).
67 Tilot, A. K. *et al.* Rare variants in axonogenesis genes connect three families with sound-color synesthesia. *Proc Natl Acad Sci U S A* **115**, 3168-3173, doi:10.1073/pnas.1715492115 (2018).
68 Boonsawat, P. *et al.* Elucidation of the phenotypic spectrum and genetic landscape in primary and secondary microcephaly. *Genet Med* **21**, 2043-2058, doi:10.1038/s41436-019-0464-7 (2019).
69 Lee, J. Y., Oh, S. H., Keum, C., Lee, B. L. & Chung, W. Y. Clinical application of prospective whole-exome sequencing in the diagnosis of genetic disease: Experience of a regional disease center in South Korea. *Ann Hum Genet* **88**, 101-112, doi:10.1111/ahg.12530 (2024).
70 Bae, H. R. & Kim, Y. O. SLC9A6-related developmental and epileptic encephalopathy with spike-and-wave activation in sleep: A case report. *Journal of Genetic Medicine* **19**, 100-104 (2022).
71 Yalcintepe, S. & Gurkan, H. Novel c.1505_1509dupCTGCC pathogenic variation in a male case with Christianson syndrome. *Clin Dysmorphol* **30**, 36-38, doi:10.1097/MCD.0000000000000358 (2021).
72 He, H. *et al.* Functional analysis of two SLC9A6 frameshift variants in lymphoblastoid cells from patients with Christianson syndrome. *CNS Neurosci Ther* **29**, 4059-4069, doi:10.1111/cns.14329 (2023).
73 Lan, Y. *et al.* Case Report: Christianson Syndrome Caused by SLC9A6 Mutation: From Case to Genotype-Phenotype Analysis. *Front Genet* **12**, 783841, doi:10.3389/fgene.2021.783841 (2021).
74 Takahashi, Y. *et al.* A loss-of-function mutation in the SLC9A6 gene causes X-linked mental retardation resembling Angelman syndrome. *Am J Med Genet B Neuropsychiatr Genet* **156B**, 799-807, doi:10.1002/ajmg.b.31221 (2011).
75 Zhou, X. *et al.* Integrating de novo and inherited variants in 42,607 autism cases identifies mutations in new moderate-risk genes. *Nat Genet* **54**, 1305-1319, doi:10.1038/s41588-022-01148-2 (2022).
76 Redin, C. *et al.* Efficient strategy for the molecular diagnosis of intellectual disability using targeted high-throughput sequencing. *J Med Genet* **51**, 724-736, doi:10.1136/jmedgenet-2014-102554 (2014).
77 Masurel-Paulet, A. *et al.* A new family with an SLC9A6 mutation expanding the phenotypic spectrum of Christianson syndrome. *Am J Med Genet A* **170**, 2103-2110, doi:10.1002/ajmg.a.37765 (2016).
78 Liu, J. *et al.* Novel and de novo mutations in pediatric refractory epilepsy. *Mol Brain* **11**, 48, doi:10.1186/s13041-018-0392-5 (2018).

79 Shu, L. *et al.* Parental mosaicism in de novo neurodevelopmental diseases. *Am J Med Genet A* **185**, 2119-2125, doi:10.1002/ajmg.a.62174 (2021).
80 Ko, A. *et al.* Targeted gene panel and genotype-phenotype correlation in children with developmental and epileptic encephalopathy. *Epilepsy Res* **141**, 48-55, doi:10.1016/j.eplepsyres.2018.02.003 (2018).
81 Trump, N. *et al.* Improving diagnosis and broadening the phenotypes in early-onset seizure and severe developmental delay disorders through gene panel analysis. *J Med Genet* **53**, 310-317, doi:10.1136/jmedgenet-2015-103263 (2016).
82 Roxrud, I., Raiborg, C., Gilfillan, G. D., Stromme, P. & Stenmark, H. Dual degradation mechanisms ensure disposal of NHE6 mutant protein associated with neurological disease. *Exp Cell Res* **315**, 3014-3027, doi:10.1016/j.yexcr.2009.07.012 (2009).
83 Ilie, A. *et al.* A Christianson syndrome-linked deletion mutation (Δ(287)ES(288)) in SLC9A6 disrupts recycling endosomal function and elicits neurodegeneration and cell death. *Mol Neurodegener* **11**, 63, doi:10.1186/s13024-016-0129-9 (2016).
84 Gao, A. Y. L., Ilie, A., Chang, P. K. Y., Orlowski, J. & McKinney, R. A. A Christianson syndrome-linked deletion mutation (Delta287ES288) in SLC9A6 impairs hippocampal neuronal plasticity. *Neurobiol Dis* **130**, 104490, doi:10.1016/j.nbd.2019.104490 (2019).
85 Zhang, X. *et al.* RT-PCR analysis of mRNA revealed the splice-altering effect of rare intronic variants in monogenic disorders. *Ann Hum Genet* **84**, 456-462, doi:10.1111/ahg.12400 (2020).
86 Slavotinek, A. *et al.* Diagnostic yield of pediatric and prenatal exome sequencing in a diverse population. *NPJ Genom Med* **8**, 10, doi:10.1038/s41525-023-00353-0 (2023).
87 Wright, J. R. *et al.* Return of genetic research results in 21,532 individuals with autism. *Genet Med* **26**, 101202, doi:10.1016/j.gim.2024.101202 (2024).
88 Garbern, J. Y. *et al.* A mutation affecting the sodium/proton exchanger, SLC9A6, causes mental retardation with tau deposition. *Brain* **133**, 1391-1402, doi:10.1093/brain/awq071 (2010).
89 Ilie, A., Weinstein, E., Boucher, A., McKinney, R. A. & Orlowski, J. Impaired posttranslational processing and trafficking of an endosomal Na+/H+ exchanger NHE6 mutant (Delta(370)WST(372)) associated with X-linked intellectual disability and autism. *Neurochem Int* **73**, 192-203, doi:10.1016/j.neuint.2013.09.020 (2014).
90 Prasad, H. & Rao, R. The Na+/H+ exchanger NHE6 modulates endosomal pH to control processing of amyloid precursor protein in a cell culture model of Alzheimer disease. *J Biol Chem* **290**, 5311-5327, doi:10.1074/jbc.M114.602219 (2015).
91 Gong, P., Xue, J., Jiao, X., Zhang, Y. & Yang, Z. Genetic Etiologies in Developmental and/or Epileptic Encephalopathy With Electrical Status Epilepticus During Sleep: Cohort Study. *Front Genet* **12**, 607965, doi:10.3389/fgene.2021.607965 (2021).
92 Yamamoto, Y. *et al.* SLC9A6-Linked Parkinson Syndrome in Female Heterozygotes Is Associated With PET-Detectable Tau Pathology. *Neurol Genet* **11**, e200235, doi:10.1212/NXG.0000000000200235 (2025).
93 Hu, H. *et al.* X-exome sequencing of 405 unresolved families identifies seven novel intellectual disability genes. *Mol Psychiatry* **21**, 133-148, doi:10.1038/mp.2014.193 (2016).
94 Zech, M. *et al.* Monogenic variants in dystonia: an exome-wide sequencing study. *Lancet Neurol* **19**, 908-918, doi:10.1016/S1474-4422(20)30312-4 (2020).
95 Dzinovic, I. *et al.* Genetic overlap between dystonia and other neurologic disorders: A study of 1,100 exomes. *Parkinsonism Relat Disord* **102**, 1-6, doi:10.1016/j.parkreldis.2022.07.003 (2022).
96 Ji, J. *et al.* A semiautomated whole-exome sequencing workflow leads to increased diagnostic yield and identification of novel candidate variants. *Cold Spring Harb Mol Case Stud* **5**, doi:10.1101/mcs.a003756 (2019).
97 Mathieu, M. L. *et al.* Electrical status epilepticus in sleep, a constitutive feature of Christianson syndrome? *Eur J Paediatr Neurol* **22**, 1124-1132, doi:10.1016/j.ejpn.2018.07.004 (2018).
98 Tzschach, A. *et al.* Christianson syndrome in a patient with an interstitial Xq26.3 deletion. *Am J Med Genet A* **155A**, 2771-2774, doi:10.1002/ajmg.a.34230 (2011).

99 Tzschach, A. *et al.* Next-generation sequencing in X-linked intellectual disability. *Eur J Hum Genet* **23**, 1513-1518, doi:10.1038/ejhg.2015.5 (2015).
100 Rim, J. H. *et al.* Efficient strategy for the molecular diagnosis of intractable early-onset epilepsy using targeted gene sequencing. *BMC Med Genomics* **11**, 6, doi:10.1186/s12920-018-0320-7 (2018).
101 Testard, Q. *et al.* Exome sequencing as a first-tier test for copy number variant detection: retrospective evaluation and prospective screening in 2418 cases. *J Med Genet* **59**, 1234-1240, doi:10.1136/jmg-2022-108439 (2022).
102 Whibley, A. C. *et al.* Fine-scale survey of X chromosome copy number variants and indels underlying intellectual disability. *Am J Hum Genet* **87**, 173-188, doi:10.1016/j.ajhg.2010.06.017 (2010).
103 Galatolo, D. *et al.* NGS in Hereditary Ataxia: When Rare Becomes Frequent. *Int J Mol Sci* **22**, doi:10.3390/ijms22168490 (2021).
104 Mantyh, P. W. *et al.* Rapid endocytosis of a G protein-coupled receptor: substance P evoked internalization of its receptor in the rat striatum in vivo. *Proc Natl Acad Sci U S A* **92**, 2622-2626, doi:10.1073/pnas.92.7.2622 (1995).
105 Gruenberg, J., Griffiths, G. & Howell, K. E. Characterization of the early endosome and putative endocytic carrier vesicles in vivo and with an assay of vesicle fusion in vitro. *J Cell Biol* **108**, 1301-1316, doi:10.1083/jcb.108.4.1301 (1989).
106 Ouyang, Q. *et al.* Christianson syndrome protein NHE6 modulates TrkB endosomal signaling required for neuronal circuit development. *Neuron* **80**, 97-112, doi:10.1016/j.neuron.2013.07.043 (2013).
107 Lee, C. *et al.* A two-domain elevator mechanism for sodium/proton antiport. *Nature* **501**, 573-577, doi:10.1038/nature12484 (2013).
108 Ishida, Y., Nayak, S., Mindell, J. A. & Grabe, M. A model of lysosomal pH regulation. *J Gen Physiol* **141**, 705-720, doi:10.1085/jgp.201210930 (2013).
109 Hartmann, T. & Verkman, A. S. Model of ion transport regulation in chloride-secreting airway epithelial cells. Integrated description of electrical, chemical, and fluorescence measurements. *Biophys J* **58**, 391-401, doi:10.1016/S0006-3495(90)82385-7 (1990).
110 Verkman, A. S. Water permeability measurement in living cells and complex tissues. *J Membr Biol* **173**, 73-87, doi:10.1007/s002320001009 (2000).
